# Separate domains of G3BP promote efficient clustering of alphavirus replication complexes and recruitment of the translation initiation machinery

**DOI:** 10.1101/600338

**Authors:** Benjamin Götte, Marc D Panas, Kirsi Hellström, Lifeng Liu, Baila Samreen, Ola Larsson, Tero Ahola, Gerald M McInerney

## Abstract

G3BP-1 and -2 (hereafter referred to as G3BP) are multifunctional RNA-binding proteins involved in stress granule (SG) assembly. Viruses from diverse families target G3BP for recruitment to replication or transcription complexes in order to block SG assembly but also to acquire pro-viral effects via other unknown functions of G3BP. The Old World alphaviruses, including Semliki Forest virus (SFV) and chikungunya virus (CHIKV) recruit G3BP into viral replication complexes, via an interaction between FGDF motifs in the C-terminus of the viral non-structural protein 3 (nsP3) and the NTF2-like domain of G3BP. To study potential proviral roles of G3BP, we used human osteosarcoma (U2OS) cell lines lacking endogenous G3BP generated using CRISPR-Cas9 and reconstituted with a panel of G3BP1 mutants and truncation variants. While SFV replicated with varying efficiency in all cell lines, CHIKV could only replicate in cells expressing G3BP1 variants containing both the NTF2-like and the RGG domains. The ability of SFV to replicate in the absence of G3BP allowed us to study effects of different domains of the protein. We used immunoprecipitation to demonstrate that that both NTF2- like and RGG domains are necessary for the formation a complex between nsP3, G3BP1 and the 40S ribosomal subunit. Electron microscopy of SFV-infected cells revealed that formation of nsP3:G3BP1 complexes via the NTF2-like domain was necessary for clustering of cytopathic vacuoles (CPVs) and that the presence of the RGG domain was necessary for accumulation of electron dense material containing G3BP1 and nsP3 surrounding the CPV clusters. Clustered CPVs also exhibited localised high levels of translation of viral mRNAs as detected by ribopuromycylation staining. These data confirm that G3BP is a ribosomal binding protein and reveal that alphaviral nsP3 uses G3BP to concentrate viral replication complexes and to recruit the translation initiation machinery, promoting the efficient translation of viral mRNAs.

## Introduction

Viral infections are inevitably accompanied by a competitive crosstalk between the host and the pathogen, engaging a complex network of protein-protein interactions. Since exploitation of host resources is crucial for the viral replication cycle, host responses aim to interfere with such measures and to clear the threat. Consequently, viruses have evolved to target host proteins involved in cellular defence mechanisms. G3BP-1 and -2 (hereafter jointly referred to as G3BP) are homologous proteins with critical roles in the assembly of cellular stress granules (SGs), dynamic assemblies of stalled translation initiation complexes and RNAs [1, 2]. The proteins contain an N-terminal NTF2-like domain, which is involved in dimerization and interaction with other proteins, long stretches of intrinsically disordered protein sequence as well as RNA-recognition motifs (RRMs) and arginine-glycine rich RGG motifs at the C terminus [2]. SG formation requires the NTF2-like and RGG domains and likely involves G3BP-driven condensation of stalled mRNP complexes as well as numerous SG nucleator proteins including TIA-1. SG induction is triggered by inhibition of translation initiation caused by cellular stresses including virus infection. As viruses strictly depend on the host translation machinery it is not surprising that many viruses from diverse virus families have developed mechanisms to disrupt SG assembly [3]. For Old World alphaviruses, including Semliki Forest virus (SFV) and chikungunya virus (CHIKV), interaction between FGDF motifs in the C-terminal region of viral non-structural protein 3 (nsP3) and G3BP disrupts SGs and blocks their formation [4, 5].

Alphaviruses are a group of enveloped positive-sense single-stranded RNA viruses belonging to the family *Togaviridae* [6]. Two separate open reading frames within the viral genome encode the non-structural and structural proteins. The four non-structural proteins, nsP1-4, form the viral replicase. Each of the non-structural proteins fulfils distinct tasks during the synthesis of viral RNA [7], but some non-structural proteins are found in other subcellular locations [8-10]. The role of nsP3 has been less well defined, but appears to be linked to several important interactions with host proteins, recently reviewed in [11] and [12]. A well-established nsP3-host interaction of the Old World alphaviruses is its binding to G3BP [4, 13-16]. This interaction is mediated by two FGDF motifs in the C-terminus of nsP3, which target a hydrophobic groove within the N-terminal NTF2-like domain of G3BP [4, 17]. In the absence of nsP3/G3BP-interactions, SFV replication is compromised, but still detectable [4, 5], while CHIKV is profoundly attenuated [16, 17]. The functional relevance of G3BP-binding during alphavirus infection has been the subject of several studies. Early events in infection lead to protein kinase R (PKR)-mediated phosphorylation of eukaryotic initiation factor 2α (eIF2α) and subsequent SG assembly [18]. G3BP is essential for SG assembly in response to eIF2α phosphorylation [2]. During the course of infection, nsP3 levels rise and SGs are disassembled through nsP3-mediated sequestration of G3BP to viral replication complexes and other locations of viral protein aggregation [4, 5, 19]. Variants of SFV mutated at both FGDF motifs, which consequently do not bind G3BP, provoke a prolonged SG response and virus growth is attenuated in cell culture [5]. Other, pro-viral roles of G3BP have been suggested, with possible functions in the switch from viral genome translation to negative-strand RNA synthesis [20] or protection of viral genomic RNA and formation of viral replication complexes [16]. In a previous study, we solved the 3-dimensional structure of the NTF2-like domain of G3BP1 (residues 1-139) in complex with a 25-residue peptide from SFV nsP3, including both FGDF motifs [17]. The NTF2-like domain forms dimers and each of the FGDF motifs binds to G3BP monomers on separate dimers, thereby crosslinking G3BP-dimers and inducing the formation of an nsP3-G3BP poly-complex. We hypothesized that this poly-complex stabilizes viral replication complexes by binding them together, producing high local concentrations of viral RNA and subsequently promoting virus replication, but functions of downstream domains of G3BP in alphavirus replication remain unidentified. Apart from the alphaviruses, many other viruses from diverse families interact with G3BP and recruit it to sites of viral macromolecule synthesis or processing [4, 21-27] but mechanisms of G3BP function in viral replication have not been described.

In this study we use a panel of G3BP1 variants to show that the NTF2-like and RGG domains of G3BP1 confer separate pro-viral functions upon recruitment to the alphaviral RNA replication complexes. The former is as a structural component of the viral RNA replication complexes, while the latter directs the recruitment of 40S ribosomal subunits to the cytopathic vacuoles (CPVs), thereby promoting efficient translation of viral mRNAs. Our work also revealed that CHIKV replication requires both these domains, while SFV replication is fully efficient with only the structural contribution of the NTF2-like domain.

## Results

### The NTF2-like and RGG domains of G3BP are necessary for efficient Old World alphavirus replication

In order to investigate the functional relevance of G3BP-recruitment to the sites of viral replication we utilised a U2OS ΔΔG3BP1/G3BP2 double-knockout cell line (hereafter referred to as U2OS ΔΔ) that was stably transfected with a selected set of different GFP-fused G3BP1 mutants (Fig. 1A), previously described in [2]. A U2OS ΔΔ cell line constitutively expressing GFP only (GFP) served as a control cell line. In addition to full-length wild type G3BP1 (GFP-G3BP WT), we used G3BP mutants or truncations that lacked specific domains or larger regions. The most extreme truncation results in expression of only the NTF2-like domain (GFP-G3BP 1-135), which preserves the ability to bind to viral nsP3 [17]. The previously described F33W mutation disrupts this interaction (GFP-G3BP F33W) [4], while maintaining SG-forming activity [2]. Additional deletions were made of distinct regions present in the C-terminal region of G3BP1, an RNA-recognition motif (GFP-G3BP ΔRRM) and an arginine/glycine-rich region (GFP-G3BP ΔRGG). G3BP mutants lacking the RGG domain fail to assemble SGs and to interact with 40S subunits [2]. The different G3BP1 variants were expressed to similar levels with exception of GFP-G3BP1 1-135, which showed variable and notably lower expression level (Fig. 1B). Densitometric analysis revealed the level to be 27% +/-8% relative to GFP-G3BP1 WT.

**Fig 1.**
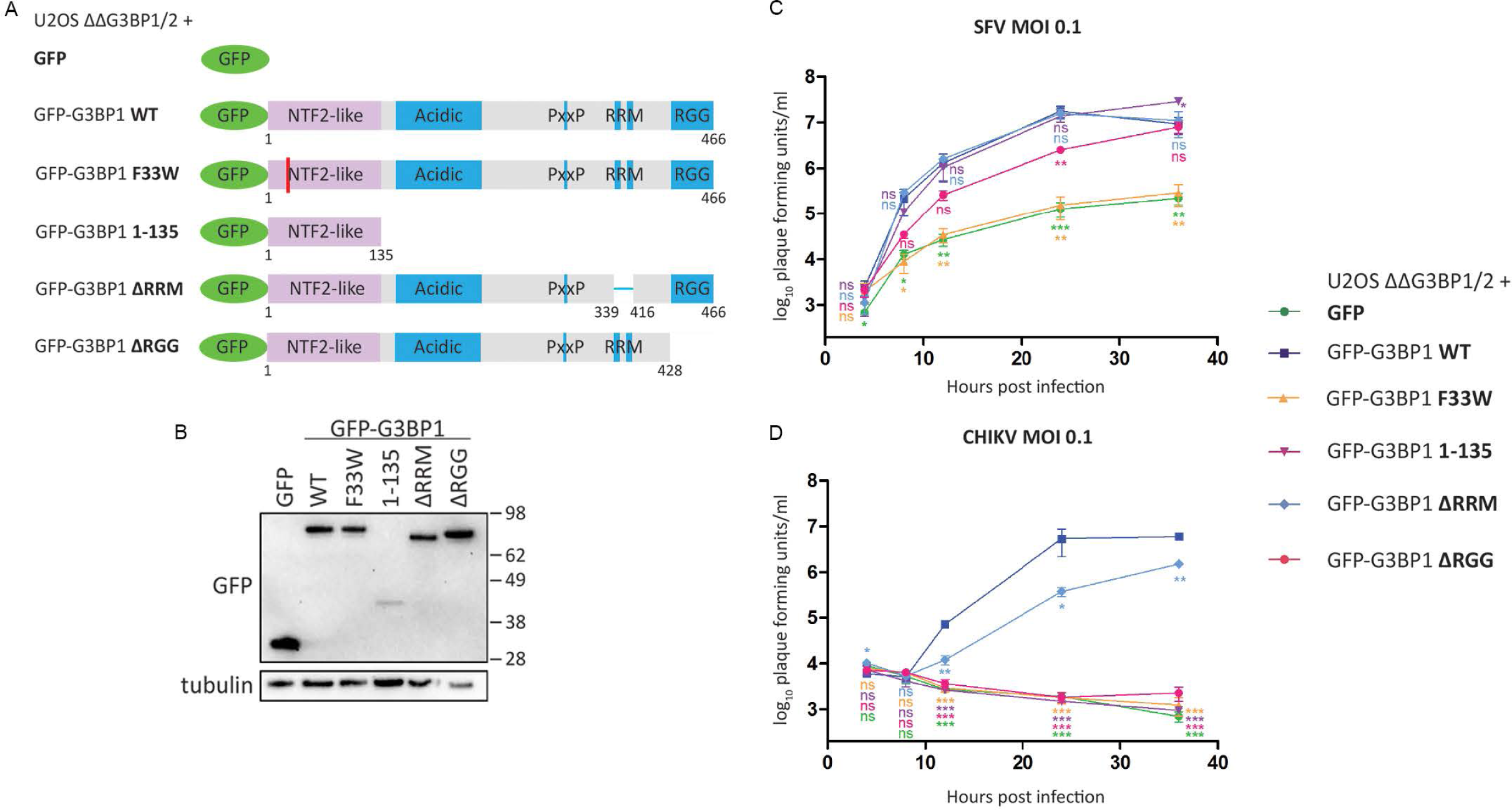
The NTF2-like and RGG domains of G3BP are necessary for efficient Old World alphavirus replication. **A.** Schematic of GFP-G3BP1 constructs used for reconstitution of G3BP1/2 double KO cell lines. **B.** Indicated cell lines were lysed and analysed by SDS-PAGE and immunoblotting for GFP and β-tubulin. **C.** and **D.** Indicated cells lines were infected with WT SFV (C) or WT CHIKV (D) at a multiplicity of infection (MOI) of 0.1. At 4, 8, 12, 24 and 36 hpi, supernatants were collected, and viral titres were quantified by plaque assay on BHK cells. Data are means of three independent experiments. Error bars indicate standard error of mean (SEM). For statistical analysis samples within each timepoint were compared to the respective titre obtained in U2OS ΔΔG3BP1/2 + GFP-G3BP1 WT cells. ns P>0.05, *P≤0.05, **P≤0.01, ***P≤0.001.

We used SFV and CHIKV as representative alphaviruses to investigate the effect of the different G3BP1 mutants on the viral life cycle in a multistep growth curve experiment. SFV replication was significantly impaired but still detectable in cells expressing GFP alone or GFP-G3BP F33W (Fig. 1C), in agreement with previous observations [4, 5]. Remarkably, expression of only the NTF2-like domain of G3BP1 (GFP-G3BP1 1-135) rescued titres similar to those obtained using full length GFP-G3BP1, despite the much lower expression of the NTF2-like domain relative to the other mutants (Fig 1B). GFP-G3BP1 ΔRRM also promoted virus replication very similarly as did GFP-G3BP1 WT, while GFP-G3BP1 ΔRGG caused a slight delay in virus growth, but still enhanced virus replication as compared to cells expressing GFP alone or GFP-G3BP F33W. Taken together, these results indicate a pro-viral effect of G3BP1 on SFV titres that is independent of G3BP’s RNA-binding, 40S subunit interaction or ability to form SGs, but is mediated by the NTF2-like domain alone.

We compared SFV replication in the GFP-G3BP1 WT cell line with the parental U2OS and the U2OS ΔΔ cells. We found that 24 hours post infection, the reconstitution rescued the viral titres to a large extent, although not completely (Fig S1A). However, at early times post infection, when we analysed the intensity of dsRNA signal per cell, we found there to be very similar levels in the parental U2OS cells as in the GFP-G3BP1 WT cell line, but barely detectable dsRNA signals in the U2OS ΔΔ cells (Fig S1B). The lack of complete rescue of SFV replication in the GFP-G3BP WT cells could be due to specific functions of G3BP2 that are lacking in the GFP-G3BP WT cells, or other effects of overexpression of the tagged protein (Fig S1C). Nevertheless, this supported that the GFP-G3BP1 WT cells are an appropriate model for studies on G3BP1 function in infection.

We next tested the growth behaviour of CHIKV in the full panel of reconstituted cell lines (Fig. 1D). In agreement with previous reports [16, 17], production of progeny virus was not detectable in absence of the nsP3-G3BP1 interaction. In contrast to SFV, the G3BP1 NTF2-like domain was unable to promote CHIKV replication. CHIKV replication was rescued by full length GFP-G3BP1 and less effectively by GFP-G3BP1 ΔRRM. Thus, CHIKV only detectably replicated in cells expressing G3BP1 variants containing both the NTF2-like and the RGG domains, suggesting that there is another G3BP-dependent proviral effect conferred on CHIKV replication by the RGG domain.

### The NTF2-like alone is required for efficient formation of SFV dsRNA-positive replication complexes

To further characterize the functional relevance of G3BP1 during the SFV and CHIKV life cycles, we performed immunofluorescence experiments to test if the different G3BP1 mutants affected the organization of viral replication complexes (RCs) within cells. For this, we first infected the cell lines presented in Fig. 1A with SFV at MOI 10, fixed them 8hpi and stained for nsP3 (red) and double stranded RNA (dsRNA, blue) (Fig. 2A). In cells expressing GFP-G3BP1 WT, the proteins colocalized with nsP3 and accumulated in close proximity to clusters of dsRNA-positive foci, indicative of sites of active viral RNA replication. In contrast, foci of nsP3 and dsRNA were detectable but weak in cells expressing GFP alone or GFP-G3BP1 F33W, and GFP signals were diffuse in the cytoplasm. In the GFP-G3BP1 F33W expressing cells, GFP signals were clearly present in large SGs in cells at early stages of infection, which were small in cells expressing high levels of nsP3. This was similar to observations made in previous studies in cells infected with SFV mutants, which were unable to bind G3BP [4, 28, 29]. Expression of the NTF2-like domain of G3BP1 (GFP-G3BP1 1-135) was sufficient to promote clustering of viral RCs in discrete foci in the cell. However, nsP3-G3BP1 1-135 aggregates appeared fewer in number and smaller in size compared to nsP3-G3BP1 WT, likely due to the lower expression level of the truncated protein. Cells expressing G3BP1 mutants lacking either the RRM (ΔRRM) or the RGG (ΔRGG) domain still produced strong dsRNA signals that colocalised with nsP3 and G3BP1, although they were less obviously clustered, and a fraction of GFP signal remained diffuse in the cytoplasm. To summarise, recruitment of G3BP1 via the NTF2-like domain is necessary for efficient formation of clusters of SFV replication complexes. This is also reflected by the quantitative analysis of dsRNA signals in images from 3 replicate experiments (Fig. 2B). Production of dsRNA-positive replication complexes is strongly impaired in the absence of the nsP3:G3BP1 interaction.

**Fig 2.**
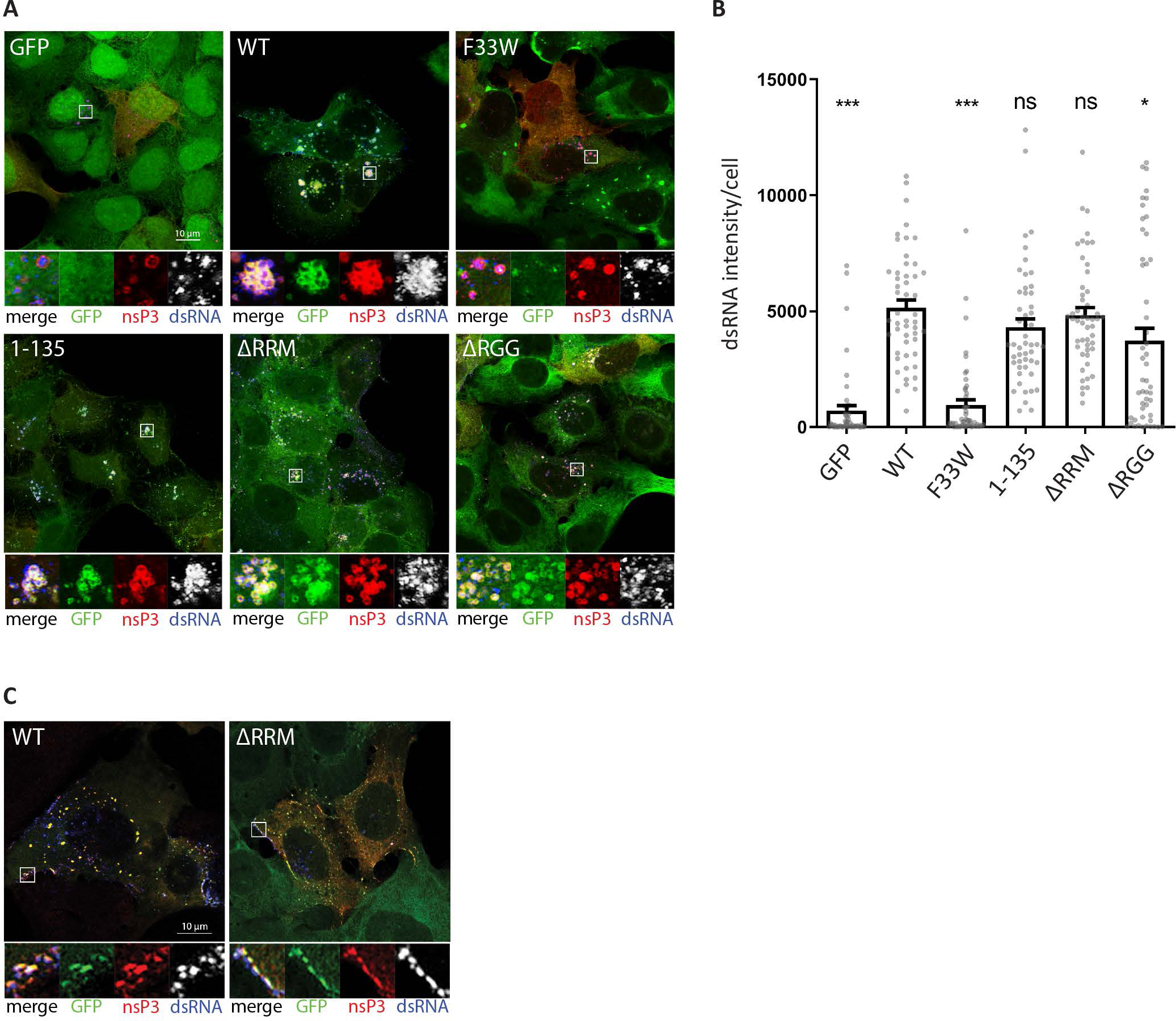
The NTF2-like and RGG domains of G3BP are differently required for efficient formation of SFV and CHIKV dsRNA-positive replication complexes. **A.** Indicated cell lines were infected with WT SFV at MOI 10. At 8 hpi cells were fixed and stained for nsP3 (red) and dsRNA (blue). Representative images from 3 independent experiments are shown. **B.** DsRNA signal intensities from images shown in A. were quantified using CellProfiler software. Bars represent mean + SEM for n = 50 cells per cell line. Individual dsRNA intensities for each analysed cell are shown as grey dots. ns P>0.05, *P≤0.05, **P≤0.01, ***P≤0.001. **C.** Indicated cell lines were infected with CHIKV at MOI 1. At 8 hpi cells were fixed and stained for nsP3 (red) and dsRNA (blue). Representative images from 3 independent experiments are shown.

### Both NTF2-like and RGG domains of G3BP1 are required for efficient formation of CHIKV dsRNA-positive replication complexes

When the same panel of cell lines was infected with CHIKV at MOI 1, only cells expressing GFP-G3BP1 WT or GFP-G3BP1 ΔRRM stained strongly for nsP3 and dsRNA (Fig. 2C), as expected from the growth curve data (Fig 1D). In GFP-G3BP1 WT cells, nsP3 was found to colocalise strongly with the GFP signal, mostly in large foci, devoid of dsRNA signal, but noticeably also colocalised with dsRNA signals in patches at the plasma membrane (Fig 2C). This was largely also true for GFP-G3BP1 ΔRRM cells although in those cultures there were fewer infected cells, containing weaker dsRNA signals and smaller nsP3-G3BP colocalizing foci. Only in very few cases (1-5% of the cells) did we observe nsP3-positive cells in GFP-G3BP1 F33W, GFP-G3BP1 1-135 or GFP-G3BP1-ΔRGG cells (Fig. S2). Moreover, in those rare cases, the dsRNA signal was either absent or at the limit of detection. Despite these differences between SFV and CHIKV, CHIKV nsP3 remained largely diffuse in GFP-G3BP1 F33W, ΔRRM and ΔRGG expressing cells, but predominantly associated with GFP-G3BP1 WT and 1-135 (Fig. 2C and S2), similar to the observations made with SFV (Fig. 2A).

### Accumulation of electron-dense patches in proximity to SFV-induced CPVs depends on the presence of the RGG domain of G3BP1

The extreme reliance of CHIKV on G3BP for viral RNA replication and gene expression makes it difficult to study the effects of mutation or truncation of the protein on viral replication. However, since SFV is less sensitive to the absence of the protein, we were able to use SFV to study the contributions of the individual domains of the protein mutated in our panel. The immunofluorescence data show that G3BP mutants influence the formation as well as organization of SFV replication complexes. To obtain a more detailed picture of this process in the different cell lines, we applied transmission electron microscopy to view infected cells. Alphavirus replication induces the formation of bulb-shaped membrane protrusions at the plasma membrane (PM), referred to as spherules, which for some viruses are later internalized to mature into cytopathic vacuoles (CPV) in the perinuclear area [30, 31]. This internalization requires another activity of nsP3, the activation of the PI3K-Akt-mTOR pathway [31, 32], found for SFV nsP3 but not CHIKV. We infected the G3BP1 mutant cell line panel with SFV and fixed at 3h and 8h post infection, to investigate the impact of GFP-G3BP1 variants on spherule formation at the PM and CPV internalization, respectively. At 3 hpi, we observed marked differences in the appearance of spherules at the PM between the different cell lines (Fig. 3A). In the absence of the nsP3-G3BP1 interaction (GFP and GFP-G3BP1 F33W cells), spherules were observed to be very few in number and sparsely distributed along the PM. In contrast, cells expressing intact G3BP1 NTF2-like domain (GFP-G3BP1 WT, 1-135, ΔRRM and ΔRGG) contained larger quantities of spherules, readily detectable at the PM. At 8 hpi (Fig. 3B), CPVs were apparent in SFV-infected GFP-G3BP1 WT, 1-135, ΔRRM and ΔRGG cells. In particular, the SFV CPVs in GFP-G3BP1 WT and 1-135 expressing cells frequently appeared in clusters of 3-5 CPVs, in some cases reaching numbers of 10 or more. As expected from the earlier time point analyses, CPVs were much less frequent in GFP and GFP-G3BP1 F33W cells, and mostly appeared isolated (Fig. 3B). From these observations, we conclude that the interaction of SFV nsP3 with the NTF2-like domain of G3BP1 is required for efficient formation of viral replication complexes early in infection and is associated with the appearance of increased numbers of CPVs, often found in clusters.

**Fig 3.**
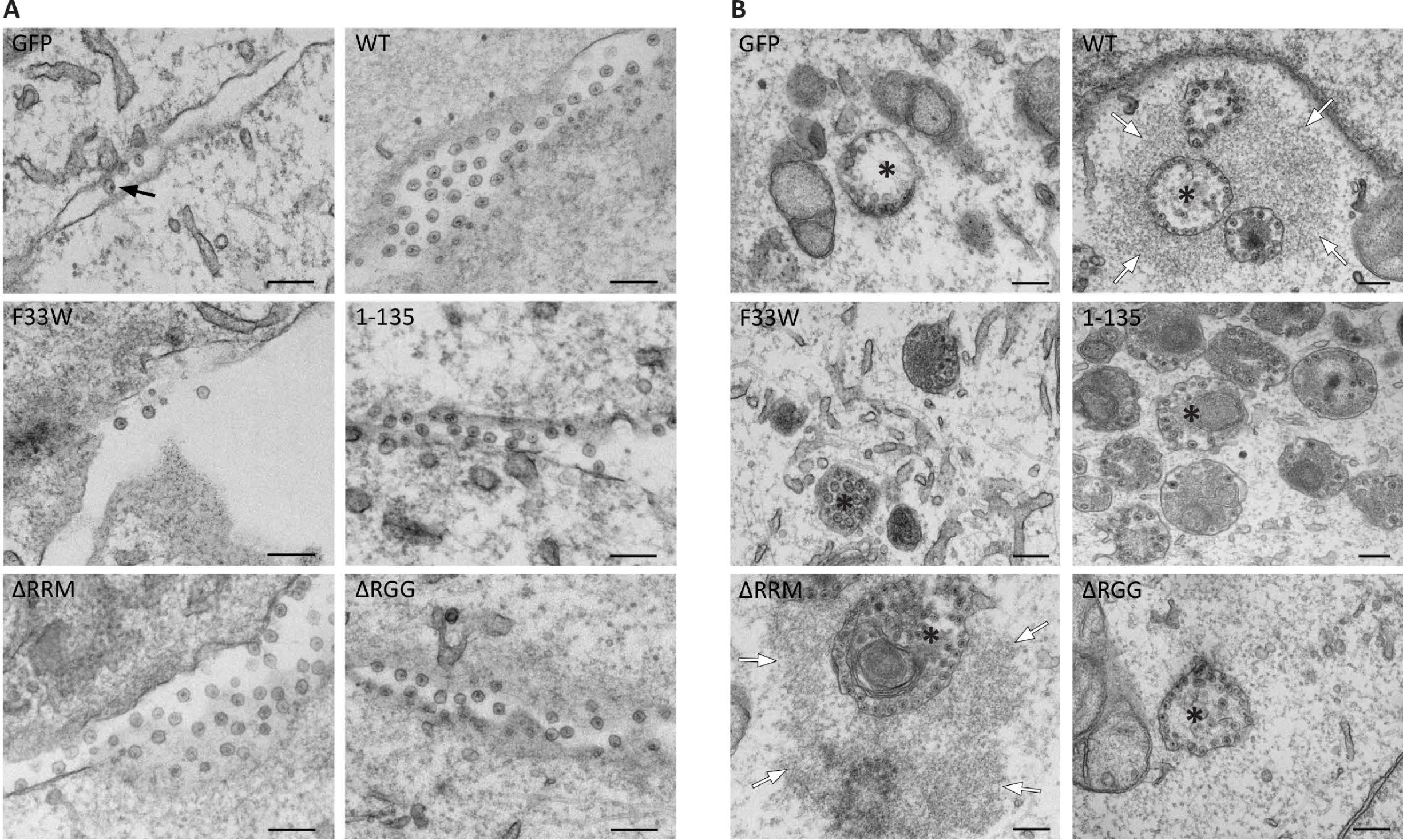
Accumulation of electron-dense patches in proximity of SFV-induced cytopathic vacuoles depend on the presence of the RGG domain of G3BP. Indicated cell lines were infected with WT SFV at MOI 100, fixed at 3 **(A)** or 8 **(B)** hpi and analysed by transmission electron microscopy. Representative images from 2 independent experiments are shown. **A.** Images of cell plasma membranes (PM) show alphavirus spherules (black arrows). **B.** Endocytosis of PM-associated spherules leads to formation of cytopathic vacuoles (CPV, asterisks) in the perinuclear region. In cells expressing GFP-G3BP1 WT and ΔRRM the CPVs are surrounded by electron-dense material (white arrows). The scale bar is 200 nm.

Although domains of G3BP downstream of the NTF2-like domain were not necessary for clustering of SFV CPVs, we noticed a distinct feature of the CPV clusters restricted to cells expressing GFP-G3BP1 WT or GFP-G3BP1 ΔRRM. In these cells, the clusters of CPVs were mostly surrounded by large (1-2 μm) patches of electron dense material, indicative of the presence of high molecular weight molecules. The formation of these patches was likely not simply a direct result of efficient RNA replication, since they were absent in SFV-infected GFP-G3BP1 1-135 cells, in which SFV replication was efficient (Fig 1C) and induced efficient clustering of CPVs, replete with spherules (Fig 3). The patches were also observed in parental U2OS cells after 8h of SFV infection (Fig S3A), excluding artefacts of GFP fusion protein overexpression, and were occasionally also detected surrounding the cytoplasmic side of spherule-containing sections of plasma membrane (Fig S3B). The CPV clusters and associated electron dense patches were frequently observed in the vicinity of membranous structures reminiscent of endoplasmic reticulum (Fig S3C). Since the GFP-G3BP1 WT and GFP-G3BP1 ΔRRM cell lines in which these patches were observed were the only lines in our panel capable of supporting the replication of CHIKV (Fig 1D), we thought it likely that these patches represent some important pro-viral factor that is required for CHIKV replication and which warranted further investigation.

These electron-dense patches bear a striking similarity to sodium arsenite-induced SGs in HeLa and HEK293 cells [33], and also to SFV infection-induced SGs which we frequently observed in U2OS ΔΔG3BP cells expressing GFP-G3BP1 F33W (Fig S3D). Overexpressed G3BP1 forms spontaneous, stress-independent SGs that require both the NTF2-like domain and RGG regions [1, 2, 34]. It is thought that the high concentrations of this protein can act to recruit ribosomes, mRNAs and other proteins containing low complexity regions and prion-like domains, which can interact via multiple low affinity interactions and undergo liquid-liquid phase separations (LLPS) to a hydrogel state [2]. The presence of the electron-dense patches around CPVs depended on the recruitment of G3BP1 via the NTF2-like domain and on the presence of the RGG region, since they were not seen in cells expressing G3BP1 variants lacking these regions. We propose therefore that the patches are generated by LLPS transitions nucleated by high local concentrations of G3BP1 surrounding clustered CPVs.

### Accumulation of GFP-G3BP1, nsP3 and translation initiation factors around SFV-induced CPVs

To identify contents of these electron-dense patches, we performed immuno-electron microscopy, staining SFV-infected U2OS ΔΔ + GFP-G3BP1 WT cells for nsP3 and GFP. In agreement with our immunofluorescence experiments, GFP-G3BP1 and nsP3 were found enriched in proximity to CPV clusters (Fig. 4A). Moreover, both proteins were not restricted to the boundaries of the CPVs, where the spherules are located, but rather stretched out over a larger area, enclosing nearby CPVs, thus likely corresponding to the patches seen in figure 3B. This was particularly noticeable for GFP-G3BP1, but to a lesser extent also for nsP3.

**Fig 4.**
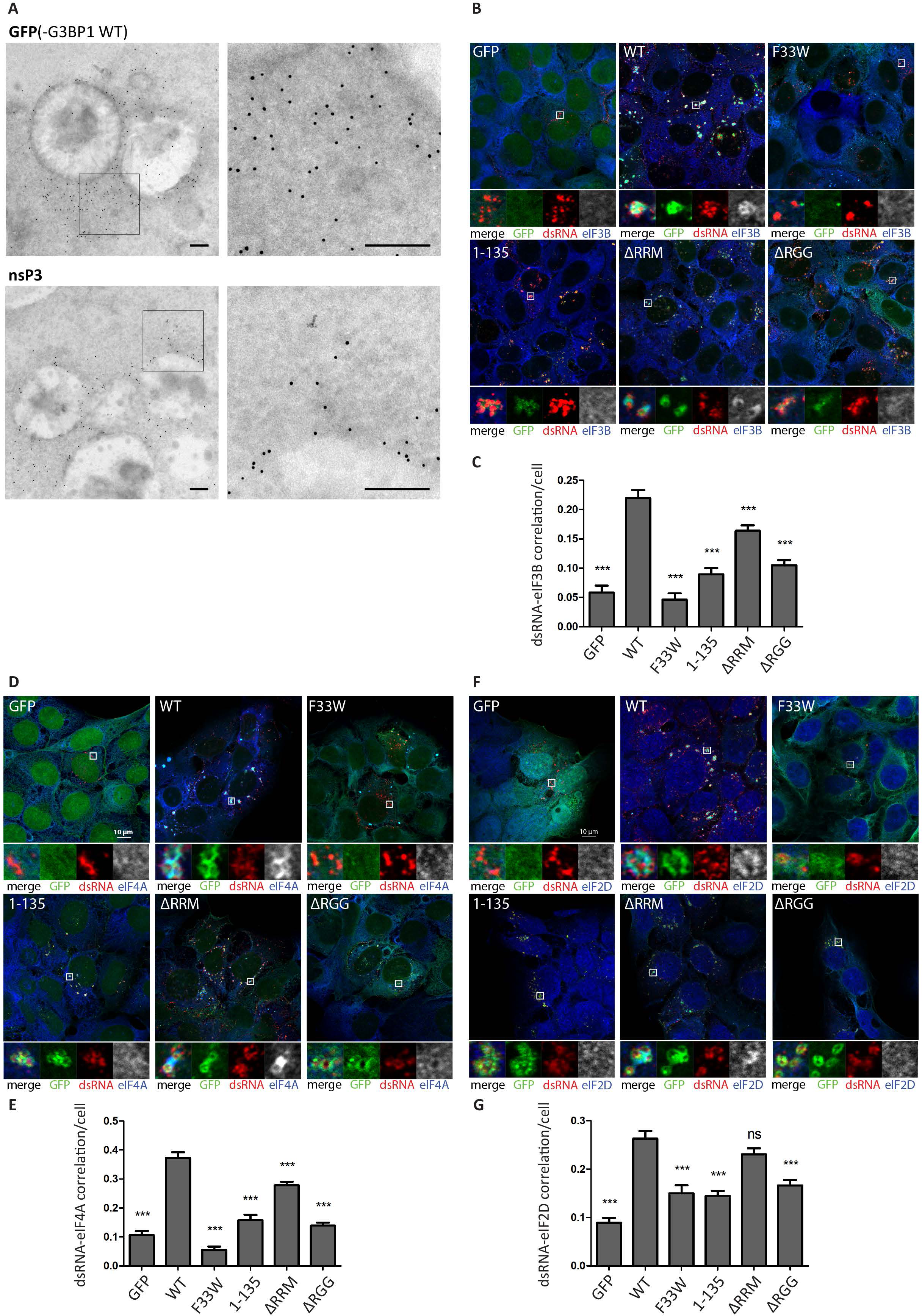
Accumulation of GFP-G3BP1, nsP3 and translation initiation factors around SFV-induced CPVs. **A.** U2OS ΔΔG3BP1/2 + GFP-G3BP1 WT cells were infected with WT SFV at MOI 100 and fixed 8h pi. Samples were prepared for immunogold labeling against GFP(-G3BP1 WT) or nsP3 and analysed by electron microscopy. Representative cytoplasmic regions that stained positive for GFP(-G3BP1 WT) or nsP3 are shown. The scale bar is 200 nm. **B.** Indicated cell lines were infected with WT SFV at MOI 10. At 8 hpi cells were fixed and stained for dsRNA (red) and **B.** eIF3B (blue) **D.** eIF4A (blue) or **F.** eIF2D (blue). Representative images from 3 independent experiments are shown. Correlations of dsRNA with eIF3B (**C.**), eIF4A (**E.**) and eIF2D (**G.**) were calculated in CellProfiler based on Pearson’s correlation coefficient for n=50 infected cells per sample. Bars represent mean + SEM. For statistical analysis indicated cell lines were compared to U2OS ΔΔG3BP1/2 + GFP-G3BP1 WT cells. ns P>0.05, *P≤0.05, **P≤0.01, ***P≤0.001.

SGs are condensates of translationally silent mRNP complexes, which include many proteins necessary for translation initiation [35, 36]. SG components are in dynamic equilibrium with polysomes, and cycle rapidly between translationally silent SGs and translationally active polysomes in cells recovering from stress [37]. We therefore considered the possibility that the recruitment of such SG components around the CPV clusters might benefit the virus, as molecules associated with translation initiation would be concentrated in the vicinity of viral mRNA production. We infected cells with SFV and stained for canonical translation initiation factors eIF3B (Fig. 4B) and eIF4A (Fig. 4D) and found that both were recruited to CPV clusters in SFV-infected GFP-G3BP1 WT-and GFP-G3BP1 ΔRRM expressing cells, but less so in those where the nsP3:G3BP1 interaction is absent or G3BP1 lacks the RGG domain. In order to quantify the degree of correlation between dsRNA and translation initiation factors, we calculated the Pearson correlation coefficient (Fig. 4C, E). For this analysis only infected cells with detectable dsRNA signals were considered. The results support the observation that the correlation is stronger for GFP-G3BP1 WT & ΔRRM cells compared to all the other cell lines, although the quantification also indicates that deletion of the RRM domain of G3BP has a negative effect on the correlation.

To verify that the recruitment of translation initiation factors is no artefact due to abnormal overexpression of GFP-G3BP1 WT, we repeated the experiment and included parental U2OS WT and U2OS ΔΔ cells. As shown in Fig. S4A, dsRNA-positive CPVs of SFV-infected U2OS WT cells strongly enrich eIF4A in close proximity, similar to U2OS ΔΔ + GFP-G3BP1 WT cells, but no such enrichment was observed in U2OS ΔΔ cells. Quantitative analysis of the degree of dsRNA-eIF4A correlation in infected dsRNA-positive cells complements this observation (Fig. S4B).

It has previously been shown that initiation of translation on the subgenomic 26S mRNA of certain alphaviruses requires the initiation factor eIF2D (previously referred to as ligatin) [38, 39]. We found that this factor is also specifically recruited to the CPV clusters in SFV-infected GFP-G3BP1 WT-and GFP-G3BP1 ΔRRM expressing cells (Fig. 4F, G).

### Old World alphavirus nsP3 forms a complex with 40S ribosomal subunits via G3BP

The RGG domain of G3BP1 is important for association with 40S subunits during formation of SGs [2]. We hypothesised therefore, that in infected cells, nsP3:G3BP1 complexes associate with 40S subunits, recruiting them to the CPVs. To determine if nsP3:G3BP:40S subunit complexes exist in SFV infected cells, we infected our panel of cell lines with SFV, isolated GFP-bound complexes at 7 hpi, and immunoblotted for nsP3 and ribosomal proteins rpS6, rpS3 and rpL4 (Fig. 5A). As expected, nsP3 strongly coprecipitated with GFP-G3BP1 WT, 1-135, ΔRRM and ΔRGG. 40S ribosomal subunit proteins rpS3 and rpS6 coprecipitated with GFP-G3BP1 WT and F33W, and even more robustly with ΔRRM, but did not detectably interact with 1-135 or ΔRGG, confirming the necessity of the RGG region for that interaction [2]. These results show that the G3BP1:40S subunit interaction remains intact in SFV-infected cells and can be found in complex with nsP3.

**Fig 5.**
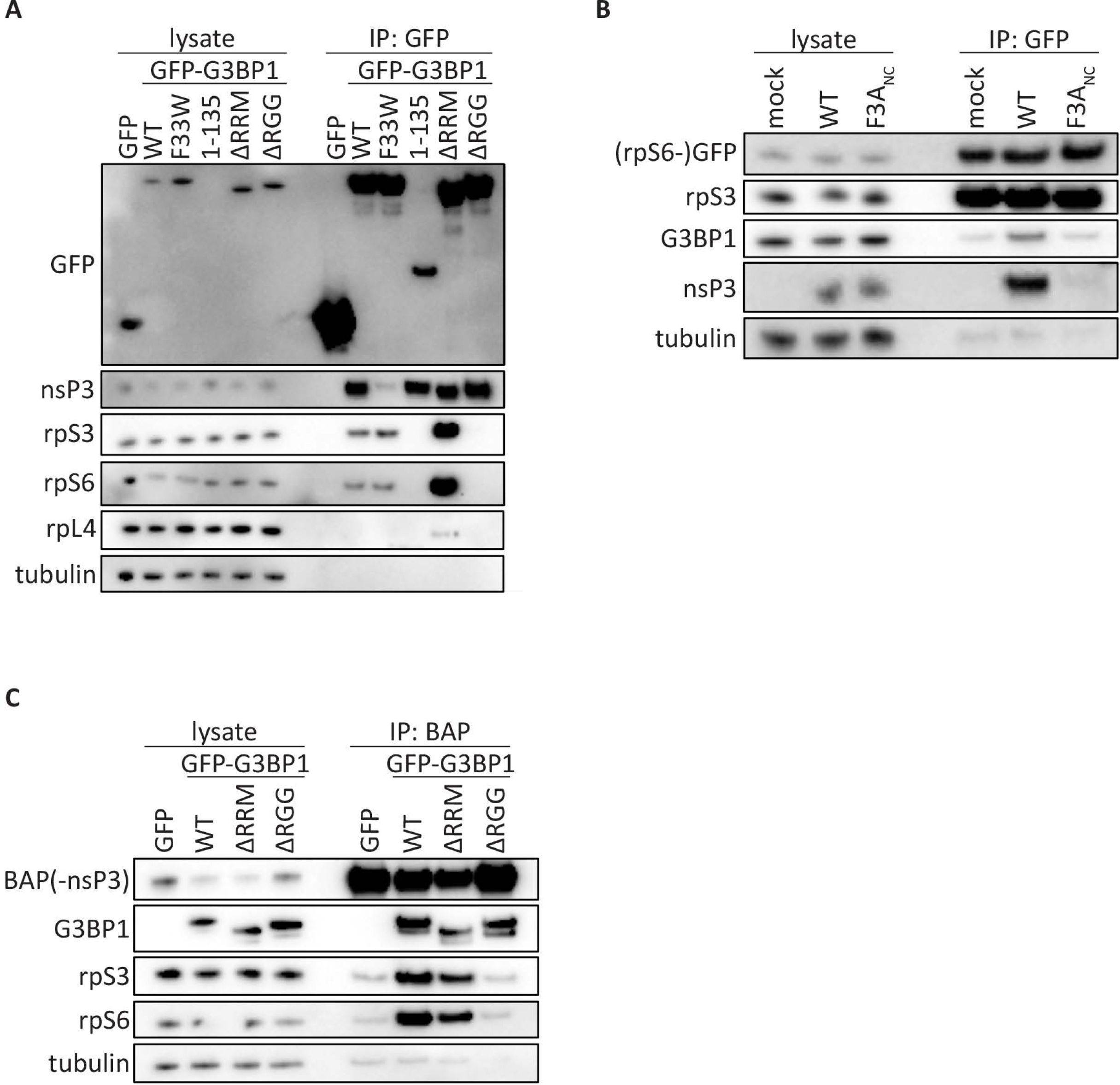
Old World alphavirus nsP3 forms a complex with 40S ribosomal subunits via G3BP. **A.** Indicated cell lines were infected with WT SFV at MOI 10. At 7 hpi, cell lysates were prepared, immunoprecipitated using GFP-Trap (ChromoTek) and separated by SDS–PAGE. Lysates and IPs were probed for GFP, nsP3, rpS3, rpS6, rpL4 or β-tubulin. **B.** GFP-rpS6 cells were infected with WT-SFV or SFV-F3A_NC_ at MOI 0.5. At 8 hpi, cell lysates were prepared, immunoprecipitated using GFP-Trap (ChromoTek) and separated by SDS–PAGE. Lysates and IPs were probed for GFP, rpS3, G3BP1, nsP3 or β-tubulin. **C.** Indicated cell lines were transfected with biotinylated CHIKV nsP3 (BAP-nsP3). Cell lysates were prepared 24 h after transfection, precipitated with streptavidin-coated beads, and separated by SDS-PAGE. Lysates and precipitates were probed with antisera for nsP3, GFP, rpS3, rpS6 or β-tubulin. All data in figure 5 are representative of at least two independent experiments.

In order to verify that ternary complexes of nsP3, G3BP and 40S subunits assemble in alphavirus infected cells, we performed immunoprecipitation experiments using pull-down of either 40S subunits or of nsP3. For pull-down of 40S ribosomal subunits, we used a U2OS cell line stably expressing rpS6-GFP, which functionally integrates into ribosomes [2]. We infected these cells with either WT SFV or SFV-F3A_NC_, which contains an inactivating point mutation in each of its two FGDF motifs, thus disrupting the nsP3:G3BP interaction [4, 17]. Isolation of 40S ribosomal subunits via rpS6-GFP was confirmed by immunostaining for rpS3. Endogenous G3BP1 coprecipitated in all conditions, but was more readily detected in SFV WT-infected cells (Fig. 5B). Importantly, nsP3 coprecipitated from SFV WT-infected cells but not from those infected with SFV-F3A_NC-_infection. This supports the observation that the nsP3:G3BP1 interaction does not prevent the association of G3BP1 with the 40S subunits, allowing the formation of nsP3:G3BP1:40S multi-protein complexes in SFV-infected cells.

The binding mechanism for CHIKV nsP3 to G3BP is very similar to that of SFV [17, 28], so we predicted that 40S subunits would also be present in CHIKV nsP3-bound complexes. We transfected a selected set of our reconstituted cell lines with biotin-acceptor peptide (BAP)-tagged CHIKV nsP3 [28, 40] and purified nsP3 complexes using streptavidin beads. Since each of the G3BP1 variants used contained an intact NTF2-like domain, all coprecipitated with nsP3, as expected (Fig. 5C). The 40S subunit proteins rpS3 and rpS6 also co-precipitated with nsP3 from lysates of GFP-G3BP1 WT and GFP-G3BP1 ΔRRM expressing cells, but not from those expressing GFP alone or GFP-G3BP1 ΔRGG.

Taken together, the results in Fig 5 demonstrate that SFV and CHIKV nsP3 form complexes with G3BP1-bound 40S ribosomal subunits in infected cells and suggest that the electron dense regions surrounding the CPV clusters contain G3BP1:40S subunit complexes as well as translation initiation factors.

### The NTF2-like and RGG domains of G3BP are required for enhanced translational activity at the CPVs

The G3BP-dependent enrichment of the translation initiation machinery in close proximity to viral replication complexes might represent a mechanism for alphaviruses to ensure efficient initiation of translation of newly produced viral RNAs. To determine whether translation of viral RNAs was more efficient when 40S subunits and initiation factors were recruited to CPVs, we used the ribopuromycylation method to visualize translational activity in our reconstituted cells after SFV infection. This method is based on the covalent attachment of puromycin (PMY) to nascent polypeptide chains produced by actively translating ribosomes and its subsequent detection using a puromycin-specific antibody [41, 42]. We infected each cell line with SFV for 8 hours, and then applied a 5-minute pulse with puromycin before fixation and staining with nsP3 and puromycin antibodies. For each cell line, representative cytoplasmic regions with characteristic nsP3-staining indicate the localization of viral CPVs (Fig. 6A, boxes). In some images, non-infected cells can be observed, containing very strong and diffuse PMY signals indicative of uninhibited translation, but in all infected cells, PMY staining is much reduced due to the profound shut off of host protein synthesis. In GFP-G3BP1 WT expressing cells, strong PMY staining was evident in and around the CPV clusters and was also detected in more diffuse areas in the cytoplasm. In contrast, GFP-G3BP1 F33W expressing cells displayed fewer clustered nsP3 puncta with no detectable PMY staining, even upon longer exposures. SFV nsP3 was also clustered in GFP-G3BP1 1-135 cells and, although PMY staining was evident close to these clusters, it was noticeably weaker than in GFP-G3BP1 WT cells, for which we had verified the interaction with 40S subunits. In GFP-G3BP1 ΔRRM expressing cells, CPVs were less clustered, but usually displayed relatively strong PMY signals. However, In GFP-G3BP1 ΔRGG expressing cells in which the G3BP:40S interaction is absent, nsP3 signals were more diffuse than in GFP-G3BP1 ΔRRM expressing cells and even when detected in clusters in some cells, contained little or no PMY staining. To allow for statistical analysis and provide a representative measure of localised translational activity, we quantified the degree of nsP3-puromycin correlation in multiple cells using the Pearson correlation coefficient (Fig. 6B). We found there to be a positive correlation in all cell lines, which was not surprising since all cells support replication and gene expression of SFV mRNA (Fig 1C). However, there was a significant reduction in correlation compared to GFP-G3BP1 WT cells in all cell lines except GFP-G3BP1 ΔRRM cells. The noticeably low levels of correlation in F33W cells may be due to the presence of SGs that are still frequently found in these cells, which would additionally restrict localized translation in these cells.

**Fig 6.**
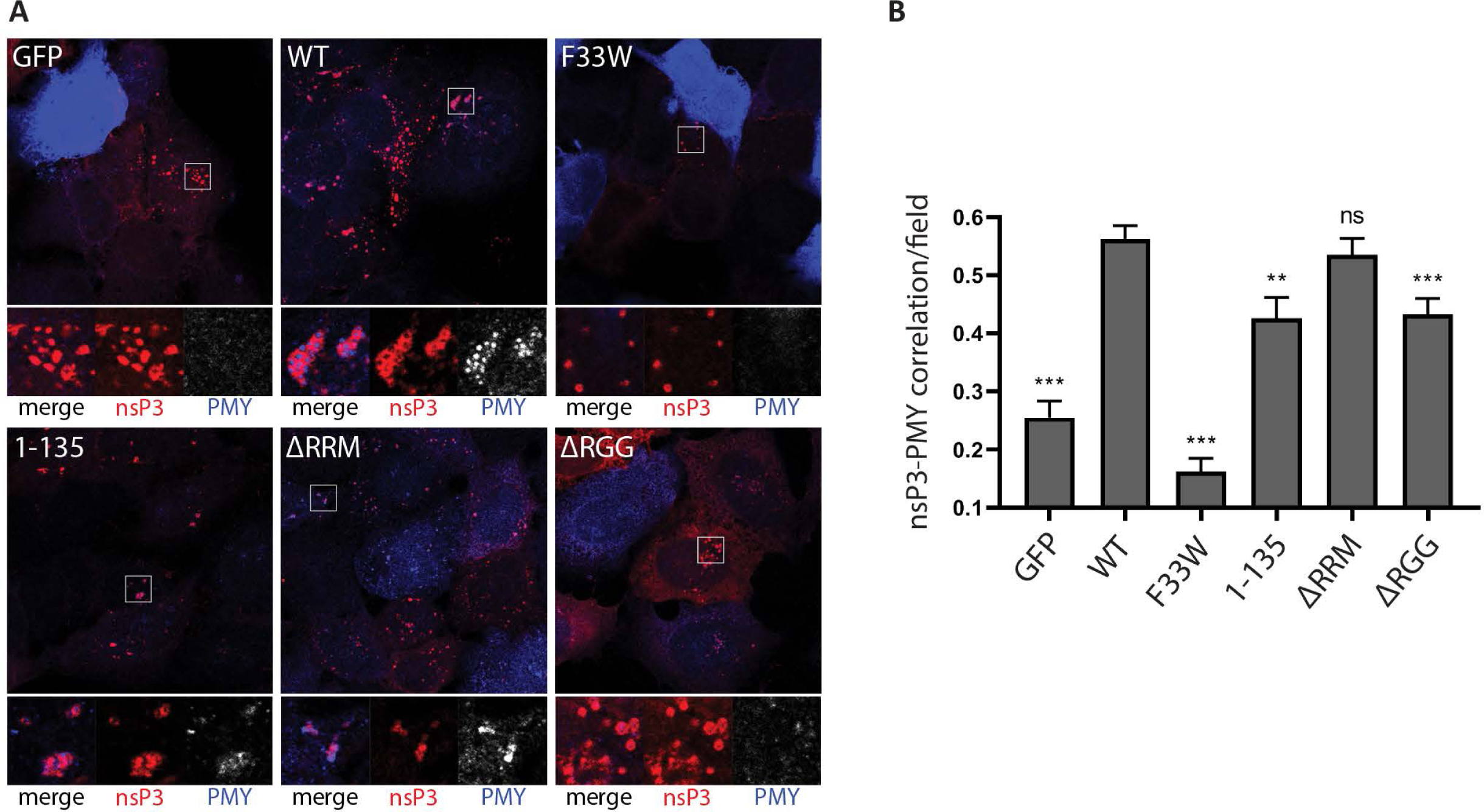
Ribopuromycylation of SFV-infected cells: The NTF2-like and RGG domains of G3BP are required for enhanced translational activity. **A.** Indicated cell lines were infected with WT SFV at MOI 10. At 8 hpi cells were pulse-labeled with puromycin (PMY) for 5 min, fixed and stained for nsP3 (red) and PMY (blue/white). Nsp3 foci indicate the localization of viral replication complexes. Staining of puromycylated nascent polypeptide chains visualizes translational activity. Representative images from 3 independent experiments are shown. **B.** Correlations of nsP3 and puromycin were calculated in CellProfiler based on Pearson’s correlation coefficient for n=20 nsP3-positive fields from ≥10 cells per cell line. Bars represent mean + SEM. For statistical analysis indicated cell lines were compared to U2OS ΔΔG3BP1/2 + GFP-G3BP1 WT cells. ns P>0.05, *P≤0.05, **P≤0.01, ***P≤0.001.

Taken together, these results support a model in which G3BP1 acts as a bridge between viral replication complexes and the 40S subunit to enhance translational activity in close proximity of viral RCs.

To further characterize the G3BP-dependent enhancement of viral mRNA translation on a cellular level, we monitored global translational efficiency using polysome preparations. This technique employs a linear sucrose gradient to separate ribosomal subunits (40S, 60S), 80S monosomes and ribosomes engaged in polysomes, and thereby allows for quantification of the relative amount of ribosomes engaged in efficient translation (i.e. associated with polysomes) [43]. We infected BHK-21 cells with either SFV WT or the G3BP-binding mutant SFV F3A_NC_ and compared their polysome tracings to mock-infected cells. Both viruses altered the polysome tracing noticeably and we observed increased absorbance at 254nM in fractions corresponding to polysomes. This is consistent with a higher number of translating ribosomes in SFV WT infected cells as compared to mock-infected cells or cells exposed to SFV F3A_NC_ (Fig. 7A, B). Thus in absence of G3BP-binding the translational efficiency in SFV-infected cells is significantly reduced. This observation is in accordance with our PMY staining, showing that recruitment of G3BP to CPVs leads to enhanced translational activity.

**Fig 7.**
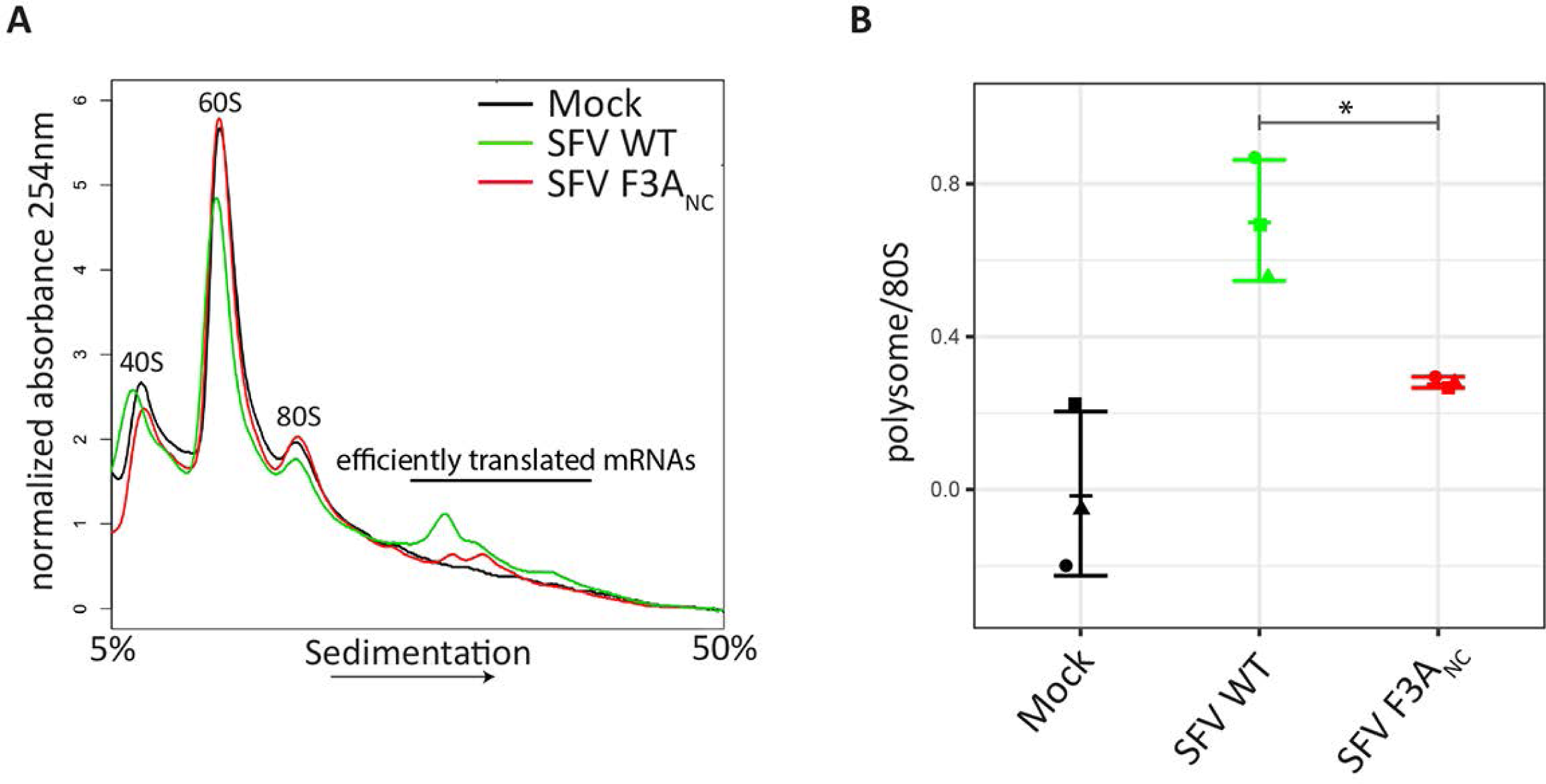
G3BP-binding enhances translation initiation in SFV-infected cells. **A.** BHK-21 cells were infected with SFV WT or SFV F3A_NC_ at MOI 10. At 8 hpi cells were treated with cycloheximide and lysates analysed on 5% - 50% sucrose gradients. Absorbance at 254 nm is shown as a function of sedimentation and fractions representing 40S, 60S, 80S and polysomes are indicated. **A** representative figure of three independent experiments is shown. **B.** The area under the curve for polysomes and the 80S peak were calculated and the ratio is shown. Data are means of three independent experiments. Error bars indicate standard deviation (SD). *P<0.05

## Discussion

The host translation machinery plays an indispensable role in viral life cycles. Despite their often very limited coding capacity, viruses have developed remarkable strategies to manipulate cellular translation to ensure synthesis of viral proteins [44]. Here we describe a novel strategy by which Old World alphaviruses utilize the cellular ribosome-associated protein G3BP1 to enrich components of the translation machinery at the sites of viral RNA replication. We show that virus replication at early stages is dependent on the NTF2-like and RGG regions of G3BP1. These are the same domains that are necessary for the formation of SGs under conditions of cellular stress, the NTF2-like domain is needed for homodimerisation as well as binding to caprin-1 and USP10, the positive and negative regulators of SGs, while the RGG region mediates binding to the 40S ribosomal subunits [2]. Under infection conditions, nsP3 binds to the N-terminal NTF2-like domain of G3BP1 and recruits it to viral replication complexes. We demonstrate that 40S ribosomal subunits remain associated with nsP3:G3BP1 complexes (Fig. 5), promoting the condensation of electron-dense patches around viral spherules at the PM and around internal CPV clusters (Figs. 3B and S3B). These electron-dense patches show noticeable similarity to bona fide SGs under the electron microscope (Fig. S3D and [33]) and are enriched for translation initiation factors that are also found in SGs [35]. In contrast to stress-induced SGs however, the nsP3:G3BP1:40S complex-dependent protein accumulations are sites of enhanced translation (Fig. 6). Our data suggest that recruitment of components of the host translation machinery to CPVs is important for the efficient synthesis of CHIKV proteins, particularly at early stages of replication and subsequently influencing all steps of the viral life cycle. This G3BP1-dependent recruitment of translational apparatus was necessary for CHIKV replication but was dispensable for SFV replication, which was as efficient in the presence of only the NTF2-like domain as it was with the full-length G3BP1 protein. Nevertheless, the translational advantage conferred by the G3BP1 RGG domain is detected in SFV infected cells. Our hypothesis is furthermore supported by the observation that infected cells, in which the SFV nsP3:G3BP1:40S complex can form (Fig 5) and engage significantly more ribosomes in translating polysomes than in cells where this complex cannot form (Fig. 7A). The sequestration of G3BP upon infection with Old World alphaviruses thus not only subverts the cellular stress response, but also more explicitly exploits a host mechanism to condense the translation machinery and target it for production of viral proteins.

The recruitment of translation initiation complexes around the CPV clusters is consistent with earlier observations of large ribonucleoprotein networks extending outwards from SFV CPVs in BHK cells that stained positive for nsP3 in immuno-EM studies [45]. In that work, the authors proposed that the networks might represent sites of production of viral RNA and proteins and of encapsidation of viral genomic RNA. Our work is consistent with that model and shows that the molecular link between the viral CPVs and the cellular translation apparatus is G3BP1, recruited by nsP3. Other, more recent work has provided evidence that the newly produced alphaviral RNAs, upon exit from the spherule neck are protected from degradation by cellular RNases [46, 47]. Indeed, a role for G3BP in protection of new viral genomic RNAs has been proposed [16], mediated by either or both of the RRM and RGG domains. Our work demonstrates a vital role for the RGG domain, and suggests that the RNAs are protected by their immediate engagement with translation initiation complexes.

RGG domains have been implicated in a range of nucleic acid and protein interactions, mediated by Arg residues in glycine-rich contexts and likely adopt a disordered and flexible structure [48]. The RGG domains of G3BP1 and 2 are classified in the Di-RG group, according to Thandapani et al [49]. Several of the Arg residues are methylated, a modification that is rapidly removed prior to SG formation [50]. It is not known whether the interaction between the G3BP1 RGG domain and the 40S ribosomal subunit is mediated by protein or RNA components of the 40S ribosome, but we previously reported that the interaction is partially resistant to RNase digestion [2]. We also note a distinct lack of aromatic amino acids in the G3BP RGG domains that would be characteristic of a nucleic acid binding domain [49]. Further work will be required to more exactly map the interaction regions of G3BP and the 40S subunit.

Our work also identified a minor role for the RRM domain, since CHIKV replication was delayed and final titres were reduced in its absence (Fig. 1D). Interestingly, we found that association of translation initiation factors with SFV dsRNA-positive replication complexes was reduced in the absence of the RRM domain (Fig. 4B-G). However, deletion of the RRM domain results in stronger association of G3BP1 with 40S subunits in GFP-G3BP1 pull-down experiments (Fig. 5A) and [2], although not in our CHIKV BAP-nsP3 immunoprecipitation (Fig. 5C). The RRM domain of G3BP is a putative RNA-binding domain containing two conserved ribonucleoprotein motifs, RNP1 and RNP2 [51]. Several studies have reported RNA-binding properties of G3BP, including interactions with viral RNA [52-54]. It is therefore possible that the RRM domain engages alphavirus mRNA once released from viral replication complexes, and thereby readily provides transcripts for 40S subunits bound to the RGG domain. Alternatively, it may be involved in the sorting of genomic RNAs away from translation complexes, for example for encapsidation. Further work will be required to determine the role of the RRM domain.

The strict requirement for particular domains of G3BP1 for CHIKV strongly restricted our experimental abilities, as CHIKV is essentially non-viable in most of our mutant cell lines (Fig. 1D). We therefore employed the closely related SFV, which replicates at a reduced level even in the absence of G3BP (Fig. 1C), in order to investigate the influence of different G3BP1 domains during various steps of the alphavirus replication cycle. SFV is commonly used as a representative model virus in Old World alphavirus research. In fact, CHIKV and SFV nsP3 proteins both contain FGDF motifs [28], which bind to the NTF2-like domain of G3BP in an identical manner although surrounding sequences may impose a slightly different orientation of the G3BP monomers in the proposed polycomplex [17]. Although the extent to which the viruses depended on G3BP1 differed, both replicated best in the presence of full length G3BP1 and least well in the absence of the protein or in the presence of the F33W binding mutant, as expected. A striking difference between CHIKV and SFV is the ability of the NTF2-like domain alone (G3BP1 1-135) to rescue the replication of SFV, but not CHIKV. The rescue of SFV replication was especially remarkable considering that the expression level of the GFP-G3BP 1-135 construct was considerably lower than that of GFP-G3BP WT (Fig. 1B). To our knowledge, no enzymatic function of the NTF2-like domain has been reported and we propose a solely structural pro-viral role. It is known that the NTF2-like domain of G3BP forms dimers [55] and in our previous work we have shown that the two FGDF-motifs of nsP3 link these dimers into a chain of nsP3-G3BP oligomers [17]. We hypothesized that these oligomeric structures could stabilize viral replication complexes by binding them together, thus ensuring that, upon internalization, each CPV contains a high concentration of spherules. In contrast to SFV, CHIKV replication was not supported by G3BP1 1-135 alone (Fig. 1D), but in the few G3BP1 1-135 expressing cells in which CHIKV nsP3 was detected, it was also observed in foci together with GFP-G3BP1 1-135 (Fig S2). However, dsRNA signals were very weak and no infectious virus was released from those cells indicating that this recruitment is not enough to promote replication of CHIKV.

Efficient rescue of CHIKV also required the presence of the RGG region and likely the recruitment of the 40S ribosomal subunits and the translation initiation factors, while for SFV this was unnecessary. What features of these two related viruses might account for the differing requirements? Compared to CHIKV, SFV generally replicates to higher titres in our various human as well as rodent cell lines. More efficient replication and consequently higher amounts of viral RNA produced might bypass the need for active recruitment of the host translation machinery. In addition, the subgenomic RNA of SFV and SINV contain a stem-loop structure, originally termed the translational enhancer, situated downstream of the initiator AUG codon, which facilitates translation initiation under conditions of low GTP–eIF2–tRNAi^Met^ ternary complex availability [18, 56, 57]. A similar RNA structure has not been detected in CHIKV subgenomic RNA [58] which might explain the increased requirement for direct recruitment of translational machinery to sites of CHIKV mRNA production. Another difference between SFV and CHIKV that could explain the need for active recruitment of the translational machinery for CHIKV is the cellular localization of viral RCs. SFV RCs are internalized from the plasma membrane (PM) and accumulate in CPV in the perinuclear area, while CHIKV RCs remain largely at the PM [31]. Naturally, exposure to the cytoplasm is more restricted for CHIKV RCs and access to translation factors in close proximity might be more limited, especially as in the initial hours of infection translation factors first accumulate in SGs, which tend to move towards the perinuclear region, away from the PM [59], unless disrupted by nsP3:G3BP interaction.

In contrast to Old World alphaviruses, the New World alphavirus VEEV is not sensitive to G3BP ablation, but efficient replication depends on another set of SG-related proteins that bind to nsP3, the Fragile X syndrome (FXR) protein family, FXR1, FXR2 and FMR1 [16]. Just like G3BP proteins, FXR proteins also associate with ribosomes, though predominantly with 60S ribosomal subunits [60]. If this interaction is important during VEEV infection remains to be investigated, but it could suggest an analogous mechanism for recruitment of ribosomes and translation initiation factors to viral replication sites by New World alphaviruses.

Targeting SG formation is a common viral strategy to interfere with inhibition of translation, often involving manipulation of G3BP [61]. The condensation of cellular translation machinery around viral RCs via sequestration of G3BP is, however, a previously unknown mechanism to ensure efficient synthesis of viral proteins. Other viruses might have evolved similar strategies, either G3BP-mediated or through other RGG domain-containing proteins, or possibly SG-associated proteins. As demonstrated here for Old World alphaviruses, this could also be a crucial step for efficient replication of other viruses, and could represent a target for antiviral therapies.

## Materials and Methods

### Cell lines and cell culture

Human osteosarcoma (U2OS) cells (ATCC HTB-96) were kept in Dulbecco’s modified Eagle’s medium (DMEM) supplemented with 10% foetal bovine serum (FBS) and 100 U/mL penicillin, and 100 µg/mL streptomycin. U2OS-derived double-null ΔΔG3BP1/2 KO cells constitutively expressing GFP-G3BP1 WT and mutants were obtained from Nancy Kedersha [2] and maintained in the same medium as wild-type U2OS supplemented with 0.5 mg/mL geneticin/G418 (Thermo Fisher Scientific). Baby hamster kidney (BHK-21) cells (ATCC CCL-10) were maintained in Glasgow’s modified Eagle’s medium (GMEM) supplemented with 10% FBS, 10% tryptose phosphate broth, 20 mM HEPES, 1mM L-glutamine and 100 U/mL penicillin, and 100 µg/mL streptomycin. All cells were cultured at 37°C in 5% CO_2_.

### Plasmids and transfection

Construction of pEBB/PP-CHIKV nsP3, encoding CHIKV nsP3 N-terminally fused to the BAP tag, is described in [40]. All plasmids were verified by sequencing (Eurofins). Cells were transiently transfected using Lipofectamine 2000 (Thermo Fisher Scientific) according to the manufacturer’s instructions.

### Viruses and virus infections

SFV was rescued by transfection of BHK-21 cells with the infectious plasmid pCMV-SFV4 [62]. Similarly, CHIKV was rescued from the pCMV-CHIKV-ICRES (CHIKV LR2006-OPY1) infectious clone. Virus titres for infection experiments on U2OS-based cell lines were determined on wild type U2OS cells by plaque assay. For infections, cell monolayers were washed with PBS and virus was added in infection medium (DMEM supplemented with 0.2% BSA, 2 mM L-glutamine, and 20 mM HEPES) with periodic shaking for 1 h at 37°C. Infectious media were then removed and cells washed with PBS before adding pre-warmed complete medium. For virus growth curve experiments, cells were grown in 6-well dishes to ∽90% confluence, infected as described above and cells overlaid with 2mL complete medium and samples of supernatant taken at different time points post infection. Virus titres were then determined on BHK cells by plaque assay.

### Virus plaque assay

A 10-fold serial dilution of virus suspension was prepared and used to infect monolayers of BHK-21 cells for 1 h at 37°C. Cells were washed with PBS, kept in BHK-21 media supplemented with 0.8% Agarose (w/v) and incubated at 37°C. After 36 h, 10% formaldehyde (v/v) in PBS was added, incubated for at least 4 h at room temperature and plaques revealed by crystal violet staining. Statistical analysis was performed using an unpaired, two-tailed Student’s t-test with a 95% confidence interval.

### SDS-PAGE and Western blotting

Samples were resolved on NuPAGE 4-12% Bis-Tris polyacrylamide gels (Thermo Fisher Scientific) and transferred onto Amersham Hybond P 0.45 PVDF blotting membranes (GE Healthcare). Membranes were blocked with 5% skim milk powder in Tris-buffered saline with 0.05% (v/v) Tween 20 (TBST) and incubated with primary antibodies (16 h at 4°C) and horseradish peroxidase (HRP)-coupled secondary antibodies (1 h at room temperature) in 1% BSA/TBST as listed below. Chemiluminescence was detected using SuperSignal West Pico PLUS Chemiluminescent Substrate (Thermo Fisher Scientific) and a Curix 60 film developer (AGFA). Primary antibodies: rabbit anti-GFP (290; Abcam; 1:10.000), mouse anti-rpS6 (74459; Santa Cruz; 1:2.000), mouse anti-rpS3 (66046-1-Ig; Proteintech; 1:2.000), rabbit antiserum against SFV-nsP3 (1:7.000; [63]), mouse anti-G3BP1 (365338; Santa Cruz; 1:1.000), goat anti-actin (1616; Santa Cruz; 1:500). Secondary antibodies: HRP-conjugated anti-mouse (Sigma A9044; 1:10.000), HRP-conjugated anti-rabbit (Cell Signaling; 1:5.000). Densitometry was performed using ImageJ.

### Immunofluorescence and microscopy

Cells grown on cover glasses (VWR) were fixed with 3.7% formaldehyde (v/v) in PBS for 15 min at room temperature, immersed in methanol for 10 min at - 20°C and blocked with 5% horse serum (Sigma) in PBS at 4°C overnight. Antibodies were diluted in blocking buffer as listed below and samples were incubated for 1 h with primary antibodies, followed by 30 min incubation with secondary antibodies at room temperature. Cover glasses were mounted on glass slides using vinol mounting media [64] and imaged by confocal laser scanning microscopy using a Supercontinuum Confocal Leica TCS SP5 X equipped with a pulsed white light laser and a Leica HCX PL Apo 63x/1.40 oil objective. Images were processed using Adobe Photoshop. Settings for image acquisition and adjustment were kept constant for all samples for dsRNA signals. Settings for nsP3 and GFP were varied slightly between samples to compensate for strong differences in localised signal intensities. Primary antibodies: mouse anti-dsRNA (English and Scientific Consulting; 1:200), rabbit antiserum against SFV-nsP3 (1:500; [63]), rabbit anti-eIF2D (Abcam ab108218; 1:200), goat anti-eIF3B (Santa Cruz Biotechnology N-20 sc-16377; 1:200), rabbit anti-eIF4A (Abcam ab31217; 1:200). Secondary antibodies: Alexa Fluor 488 (Molecular Probes; 1:200), Alexa Fluor 568 (Molecular Probes; 1:1.000), Alexa Fluor 647 (Molecular Probes; 1:500).

### Image analysis

Immunofluorescence images were analysed in CellProfiler [65]. Settings for confocal image acquisition were kept constant among all cell lines to ensure comparability during the analysis procedure. In CellProfiler, cells were detected using the ‘IdentifyPrimaryObjects’, ‘IdentifySecondaryObjects’ and ‘IdentifyTertiaryObjects’ modules. For each identified cell object, dsRNA signals were quantified using the ‘MeasureObjectIntensity’ module and correlations between dsRNA and initiation factors (eIF2D, eIF3B, eIF4A) were calculated as Pearson’s correlation coefficient using the ‘MeasureColocalization’ module. Calculations of correlation were performed on infected cells only, based on the dsRNA signal. Correlations between nsP3 and puromycin were performed on selected nsP3-positive fields of fixed size. Statistical analysis was done using an unpaired, two-tailed Student’s t-test with a 95% confidence interval.

### Immunoprecipitation

Near-confluent cells grown in a 100-mm dish were washed with cold phosphate-buffered saline without calcium and magnesium (PBS(-)) and scrape-harvested in 600µl cold EE-buffer (50 mM HEPES, pH 7.0, 150 mM NaCl, 0.1% NP-40, 10% glycerol, 4 mM EDTA, 2.5 mM EGTA, 0.1 mg/mL Heparin (H3149, Sigma), 1 mM DTT, and HALT protease inhibitors (Thermo Fisher Scientific))[2]. Cells were sonicated twice for 2 min in an ice-water bath (BRANSON 1510), rotated for 15 min at 4°C and cleared by centrifugation (10.000 g, 10 min, 4°C). GFP-fusion protein-complexes were immunoprecipitated with 25 µl GFP-Trap (ChromoTek) for 60 min at 4°C under constant rotation. Beads were then washed three times with cold EE-buffer and eluted into 2x NuPAGE LDS sample buffer (Thermo Fisher Scientific) containing 50 mM DTT, heated at 80°C for 10 min and analysed by SDS-PAGE and western blotting.

### Ribopuromycylation assay

Ribopuromycylation was modified from [42], as described in [41]. In brief, cells were left uninfected or infected with SFV WT at a MOI of 10 for 8h. 5 min before fixation, puromycin was diluted in 37°C prewarmed complete medium to a final concentration of 10.6 µM, added to the cells and incubated for 5 min at 37°C, 5% CO2. Cells were then washed with PBS and fixed with 3.7% formaldehyde (Sigma) for 5 min and permeabilized for 5 min with ice-cold methanol (Sigma). Cells were then washed with PBS and blocked with 5% horse serum for 30 min at RT. After blocking cells were stained with an anti-puromycin antibody (MABE343; Millipore; 1:1,000) and anti-SFV nsP3 antibody (1:500; [63]) for 2h at RT. Cells without puromycin treatment were used as negative controls. Images were taken with a Supercontinuum Confocal Leica TCS SP5 X equipped with a pulsed white light laser and a Leica HCX PL Apo 63x/1.40 Oil objective and processed using Adobe Photoshop. Settings for image acquisition and adjustment were kept constant for all samples.

### Transmission electron microscopy

Cells were grown on no.1 cover glasses (VWR) and infected with SFV4 at a multiplicity of infection (MOI) of 100. At 3 h and 8 h post infection (pi), cells were fixed with 2% glutaraldehyde in 0.1M sodium cacodylate buffer for 30 min at room temperature in the dark. After fixation, cells were washed three times with 0.1M sodium cacodylate buffer, stained with reduced buffered osmium tetroxide and uranyl acetate and processed for flat embedding and ultrathin sectioning as described in [67]. Images were taken with a Jeol JEM-1400 microscope (80 kV) and a bottom-mounted camera, Gatan Orius SC 1000B.

### Immunoelectron microscopy

Cells were fixed in 3 % paraformaldehyde in 0.1 M phosphate buffer (PB). Samples were rinsed in 0.1 M PB and infiltrated in 10% gelatin. Specimens were then infiltrated into 2.3 M of sucrose and frozen in liquid nitrogen. Sectioning was performed at −95°C and placed on carbon-reinforced formvar-coated, 50 mesh Nickel grids. Immunolabelling procedure was performed as follows: grids were placed directly on drops of 2% bovine serum albumin (BSA) + 2% gelatin in 0.1 M PB for 1 hour to block non-specific binding. Sections were then incubated with the primary antibody diluted in 0.1 M of phosphate buffer containing 0.2% BSA and 0.2% gelatin overnight in a humidified chamber at room temperature. The sections were thoroughly washed in the same buffer and bound antibodies were detected with protein A coated with 10 nm gold (BBI solution, Analytic standard, Sweden) at a final dilution of 1:100. Sections were rinsed in buffer and fixed in 2% glutaraldehyde and contrasted with 0.05% uranyl acetate and embedded in 1% methylcellulose and examined in a examined in a Tecnai G2 Bio TWIN (FEI company, Eindhoven, The Netherlands) at 100 kV. Digital images were taken using a Veleta camera (Soft Imaging System GmbH, Munster, Germany) [68].

### Polysome profiling

BHK cells were seeded in 15cm plates and cultured for 48h until 60-70% confluence. Following an 8h infection, polysome fractionation was performed as previously described [66] and the absorbance at 254 nm was recorded along a 5-50% sucrose gradient. The polysome-tracings were normalized to the total absorbance signal (i.e. the area under the curve for the 60S ribosomal subunit, the 80S ribosome and polysomes). Global translational efficiency was quantified as the ratio of polysomes (area under the curve of polysomes) over 80S ribosomes (area under the curve of 80S ribosomes).

## Acknowledgements

The authors wish to thank Juliane Brun for technical help and Leo Hanke and Nancy Kedersha for critical reading of the manuscript. BG gratefully acknowledges a studentship from Karolinska Institutet (KI Doktorander).

## Figure Legends

**Supporting Fig S1. GFP-G3BP1 WT rescues SFV replication.** Indicated cells lines were infected with WT SFV at a MOI of 0.1. At 24 hpi, supernatants were collected, and viral titres were quantified by plaque assay on BHK cells. Data are means of three independent experiments. Error bars indicate standard deviation (SD). **B.** Indicated cell lines were infected with WT SFV at MOI 10. At 8 hpi cells were fixed, stained for dsRNA, and dsRNA signal intensities per cell were quantified using CellProfiler software. Bars represent mean + SEM for n = 50 cells per cell line. Individual dsRNA intensities for each analysed cell are shown as grey dots. For statistical analysis indicated cell lines were compared to U2OS WT cells or as indicated. ns P>0.05, *P≤0.05, **P≤0.01, ***P≤0.001. **C.** U2OS WT or U2OS ΔΔ + GFP-G3BP1 WT cells were lysed and analysed by SDS-PAGE and immunoblotting for G3BP1 and actin.

**Supporting Fig S2. CHIKV infection depends on the NTF2-like and RGG domains of G3BP.** Indicated cell lines were infected with CHIKV at MOI 1. At 8 hpi cells were fixed and stained for nsP3 (red) and dsRNA (blue). Representative images of rare dsRNA-positive cells from 3 independent experiments are shown.

**Supporting Fig S3. Electron-dense patches surround SFV-induced cytopathic vacuoles. A.** U2OS WT cells. **B.** and **C.** U2OS ΔΔ + GFP-G3BP1 WT and **D.** U2OS ΔΔ + GFP-G3BP1 F33W cell lines were infected with WT SFV at MOI 100, fixed at 8 hpi and analysed by transmission electron microscopy.. Black arrows, spherules; White arrows, electron-dense material; Asterisks, cytopathic vacuoles. The scale bar is 200 nm.

**Supporting Fig S4. Recruitment of eIF4A to SFV CPVs in parental U2OS cells. A.** Indicated cell lines were infected with WT SFV at MOI 10. At 8 hpi cells were fixed and stained for dsRNA (red) and eIF4A (blue/white). Representative images from 2 independent experiments are shown. **B.** Correlation of dsRNA with eIF4A was calculated in CellProfiler based on Pearson’s correlation coefficient for n=50 infected cells per sample. Bars represent mean + SEM. For statistical analysis indicated cell lines were compared to U2OS WT cells or as indicated. **P≤0.01, ***P≤0.001.

## Bibliography

1. Tourriere H, Chebli K, Zekri L, Courselaud B, Blanchard JM, Bertrand E, et al. The RasGAP-associated endoribonuclease G3BP assembles stress granules. J Cell Biol. 2003;160(6):823–31. Epub 2003/03/19. doi: 10.1083/jcb.200212128. PubMed PMID: 12642610; PubMed Central PMCID: PMCPMC2173781.

2. Kedersha N, Panas MD, Achorn CA, Lyons S, Tisdale S, Hickman T, et al. G3BP-Caprin1-USP10 complexes mediate stress granule condensation and associate with 40S subunits. J Cell Biol. 2016;212(7):845–60. Epub 2016/03/30. doi: 10.1083/jcb.201508028. PubMed PMID: 27022092; PubMed Central PMCID: PMCPMC4810302.

3. Reineke LC, Lloyd RE. Diversion of stress granules and P-bodies during viral infection. Virology. 2013;436(2):255–67. Epub 2013/01/08. doi: 10.1016/j.virol.2012.11.017. PubMed PMID: 23290869; PubMed Central PMCID: PMCPMC3611887.

4. Panas MD, Schulte T, Thaa B, Sandalova T, Kedersha N, Achour A, et al. Viral and cellular proteins containing FGDF motifs bind G3BP to block stress granule formation. PLoS Pathog. 2015;11(2):e1004659. Epub 2015/02/07. doi: 10.1371/journal.ppat.1004659. PubMed PMID: 25658430; PubMed Central PMCID: PMCPMC4450067.

5. Panas MD, Varjak M, Lulla A, Eng KE, Merits A, Karlsson Hedestam GB, et al. Sequestration of G3BP coupled with efficient translation inhibits stress granules in Semliki Forest virus infection. Mol Biol Cell. 2012;23(24):4701–12. Epub 2012/10/23. doi: 10.1091/mbc.E12-08-0619. PubMed PMID: 23087212; PubMed Central PMCID: PMCPMC3521679.

6. Chen R, Mukhopadhyay S, Merits A, Bolling B, Nasar F, Coffey LL, et al. ICTV Virus Taxonomy Profile: Togaviridae. J Gen Virol. 2018;99(6):761–2. Epub 2018/05/11. doi: 10.1099/jgv.0.001072. PubMed PMID: 29745869.

7. Rupp JC, Sokoloski KJ, Gebhart NN, Hardy RW. Alphavirus RNA synthesis and non-structural protein functions. J Gen Virol. 2015;96(9):2483–500. Epub 2015/07/30. doi: 10.1099/jgv.0.000249. PubMed PMID: 26219641; PubMed Central PMCID: PMCPMC4635493.

8. Gorchakov R, Garmashova N, Frolova E, Frolov I. Different types of nsP3-containing protein complexes in Sindbis virus-infected cells. J Virol. 2008;82(20):10088–101. Epub 2008/08/08. doi: 10.1128/JVI.01011-08. PubMed PMID: 18684830; PubMed Central PMCID: PMCPMC2566286.

9. Peranen J, Laakkonen P, Hyvonen M, Kaariainen L. The alphavirus replicase protein nsP1 is membrane-associated and has affinity to endocytic organelles. Virology. 1995;208(2):610–20. doi: 10.1006/viro.1995.1192. PubMed PMID: 7747433.

10. Peranen J, Rikkonen M, Liljestrom P, Kaariainen L. Nuclear localization of Semliki Forest virus-specific nonstructural protein nsP2. J Virol. 1990;64(5):1888–96. PubMed PMID: 2139138.

11. Gotte B, Liu L, McInerney GM. The Enigmatic Alphavirus Non-Structural Protein 3 (nsP3) Revealing Its Secrets at Last. Viruses. 2018;10(3). Epub 2018/03/03. doi: 10.3390/v10030105. PubMed PMID: 29495654; PubMed Central PMCID: PMCPMC5869498.

12. Lark T, Keck F, Narayanan A. Interactions of Alphavirus nsP3 Protein with Host Proteins. Front Microbiol. 2017;8:2652. Epub 2018/01/30. doi: 10.3389/fmicb.2017.02652. PubMed PMID: 29375517; PubMed Central PMCID: PMCPMC5767282.

13. Cristea IM, Carroll JW, Rout MP, Rice CM, Chait BT, MacDonald MR. Tracking and elucidating alphavirus-host protein interactions. J Biol Chem. 2006;281(40):30269–78. Epub 2006/08/10. doi: 10.1074/jbc.M603980200. PubMed PMID: 16895903.

14. Frolova E, Gorchakov R, Garmashova N, Atasheva S, Vergara LA, Frolov I. Formation of nsP3-specific protein complexes during Sindbis virus replication. J Virol. 2006;80(8):4122–34. Epub 2006/03/31. doi: 10.1128/JVI.80.8.4122-4134.2006. PubMed PMID: 16571828; PubMed Central PMCID: PMCPMC1440443.

15. Frolov I, Kim DY, Akhrymuk M, Mobley JA, Frolova EI. Hypervariable Domain of Eastern Equine Encephalitis Virus nsP3 Redundantly Utilizes Multiple Cellular Proteins for Replication Complex Assembly. J Virol. 2017;91(14). Epub 2017/05/05. doi: 10.1128/JVI.00371-17. PubMed PMID: 28468889; PubMed Central PMCID: PMCPMC5487569.

16. Kim DY, Reynaud JM, Rasalouskaya A, Akhrymuk I, Mobley JA, Frolov I, et al. New World and Old World Alphaviruses Have Evolved to Exploit Different Components of Stress Granules, FXR and G3BP Proteins, for Assembly of Viral Replication Complexes. PLoS Pathog. 2016;12(8):e1005810. Epub 2016/08/11. doi: 10.1371/journal.ppat.1005810. PubMed PMID: 27509095; PubMed Central PMCID: PMCPMC4980055.

17. Schulte T, Liu L, Panas MD, Thaa B, Dickson N, Gotte B, et al. Combined structural, biochemical and cellular evidence demonstrates that both FGDF motifs in alphavirus nsP3 are required for efficient replication. Open Biol. 2016;6(7). Epub 2016/07/08. doi: 10.1098/rsob.160078. PubMed PMID: 27383630; PubMed Central PMCID: PMCPMC4967826.

18. McInerney GM, Kedersha NL, Kaufman RJ, Anderson P, Liljestrom P. Importance of eIF2alpha phosphorylation and stress granule assembly in alphavirus translation regulation. Mol Biol Cell. 2005;16(8):3753–63. Epub 2005/06/03. doi: 10.1091/mbc.E05-02-0124. PubMed PMID: 15930128; PubMed Central PMCID: PMCPMC1182313.

19. Fros JJ, Domeradzka NE, Baggen J, Geertsema C, Flipse J, Vlak JM, et al. Chikungunya virus nsP3 blocks stress granule assembly by recruitment of G3BP into cytoplasmic foci. J Virol. 2012;86(19):10873–9. Epub 2012/07/28. doi: 10.1128/JVI.01506-12. PubMed PMID: 22837213; PubMed Central PMCID: PMCPMC3457282.

20. Scholte FE, Tas A, Albulescu IC, Zusinaite E, Merits A, Snijder EJ, et al. Stress granule components G3BP1 and G3BP2 play a proviral role early in Chikungunya virus replication. J Virol. 2015;89(8):4457–69. Epub 2015/02/06. doi: 10.1128/JVI.03612-14. PubMed PMID: 25653451; PubMed Central PMCID: PMCPMC4442398.

21. Katsafanas GC, Moss B. Colocalization of transcription and translation within cytoplasmic poxvirus factories coordinates viral expression and subjugates host functions. Cell Host Microbe. 2007;2(4):221–8. Epub 2007/11/17. doi: S1931-3128(07)00215-6 [pii] 10.1016/j.chom.2007.08.005. PubMed PMID: 18005740; PubMed Central PMCID: PMC2084088.

22. Krapp S, Greiner E, Amin B, Sonnewald U, Krenz B. The stress granule component G3BP is a novel interaction partner for the nuclear shuttle proteins of the nanovirus pea necrotic yellow dwarf virus and geminivirus abutilon mosaic virus. Virus Res. 2017;227:6–14. doi: 10.1016/j.virusres.2016.09.021. PubMed PMID: 27693920.

23. Tsai KN, Chong CL, Chou YC, Huang CC, Wang YL, Wang SW, et al. Doubly Spliced RNA of Hepatitis B Virus Suppresses Viral Transcription via TATA-Binding Protein and Induces Stress Granule Assembly. J Virol. 2015;89(22):11406–19. doi: 10.1128/JVI.00949-15. PubMed PMID: 26339052; PubMed Central PMCID: PMCPMC4645676.

24. Matthews JD, Frey TK. Analysis of subcellular G3BP redistribution during rubella virus infection. J Gen Virol. 2012;93(Pt 2):267–74. doi: 10.1099/vir.0.036780-0. PubMed PMID: 21994324; PubMed Central PMCID: PMCPMC3352340.

25. Simpson-Holley M, Kedersha N, Dower K, Rubins KH, Anderson P, Hensley LE, et al. Formation of antiviral cytoplasmic granules during orthopoxvirus infection. J Virol. 2011;85(4):1581–93. Epub 2010/12/15. doi: JVI.02247-10 [pii] 10.1128/JVI.02247-10. PubMed PMID: 21147913; PubMed Central PMCID: PMC3028896.

26. Ariumi Y, Kuroki M, Kushima Y, Osugi K, Hijikata M, Maki M, et al. Hepatitis C virus hijacks P-body and stress granule components around lipid droplets. J Virol. 2011;85(14):6882–92. doi: 10.1128/JVI.02418-10. PubMed PMID: 21543503; PubMed Central PMCID: PMCPMC3126564.

27. Pager CT, Schutz S, Abraham TM, Luo G, Sarnow P. Modulation of hepatitis C virus RNA abundance and virus release by dispersion of processing bodies and enrichment of stress granules. Virology. 2013;435(2):472–84. Epub 2012/11/13. doi: S0042-6822(12)00535-1 [pii] 10.1016/j.virol.2012.10.027. PubMed PMID: 23141719; PubMed Central PMCID: PMC3534916.

28. Panas MD, Ahola T, McInerney GM. The C-terminal repeat domains of nsP3 from the Old World alphaviruses bind directly to G3BP. J Virol. 2014;88(10):5888–93. Epub 2014/03/14. doi: 10.1128/JVI.00439-14. PubMed PMID: 24623412; PubMed Central PMCID: PMCPMC4019107.

29. Panas MD, Varjak M, Lulla A, Er Eng K, Merits A, Karlsson Hedestam GB, et al. Sequestration of G3BP coupled with efficient translation inhibits stress granules in Semliki Forest virus infection. Mol Biol Cell. 2012;23(24):4701–12. Epub 2012/10/23. doi: mbc.E12-08-0619 [pii] 10.1091/mbc.E12-08-0619. PubMed PMID: 23087212; PubMed Central PMCID: PMC3521679.

30. Spuul P, Balistreri G, Kaariainen L, Ahola T. Phosphatidylinositol 3-kinase-, actin-, and microtubule-dependent transport of Semliki Forest Virus replication complexes from the plasma membrane to modified lysosomes. J Virol. 2010;84(15):7543–57. Epub 2010/05/21. doi: 10.1128/JVI.00477-10. PubMed PMID: 20484502; PubMed Central PMCID: PMCPMC2897599.

31. Thaa B, Biasiotto R, Eng K, Neuvonen M, Gotte B, Rheinemann L, et al. Differential Phosphatidylinositol-3-Kinase-Akt-mTOR Activation by Semliki Forest and Chikungunya Viruses Is Dependent on nsP3 and Connected to Replication Complex Internalization. J Virol. 2015;89(22):11420–37. Epub 2015/09/05. doi: 10.1128/JVI.01579-15. PubMed PMID: 26339054; PubMed Central PMCID: PMCPMC4645633.

32. Mazzon M, Castro C, Thaa B, Liu L, Mutso M, Liu X, et al. Alphavirus-induced hyperactivation of PI3K/AKT directs pro-viral metabolic changes. PLoS Pathog. 2018;14(1):e1006835. doi: 10.1371/journal.ppat.1006835. PubMed PMID: 29377936.

33. Souquere S, Mollet S, Kress M, Dautry F, Pierron G, Weil D. Unravelling the ultrastructure of stress granules and associated P-bodies in human cells. J Cell Sci. 2009;122(Pt 20):3619–26. Epub 2009/10/09. doi: 10.1242/jcs.054437. PubMed PMID: 19812307.

34. Matsuki H, Takahashi M, Higuchi M, Makokha GN, Oie M, Fujii M. Both G3BP1 and G3BP2 contribute to stress granule formation. Genes Cells. 2013;18(2):135–46. Epub 2013/01/03. doi: 10.1111/gtc.12023. PubMed PMID: 23279204.

35. Kedersha N, Chen S, Gilks N, Li W, Miller IJ, Stahl J, et al. Evidence that ternary complex (eIF2-GTP-tRNA(i)(Met))-deficient preinitiation complexes are core constituents of mammalian stress granules. Mol Biol Cell. 2002;13(1):195–210. PubMed PMID: 11809833.

36. Kedersha NL, Gupta M, Li W, Miller I, Anderson P. RNA-binding proteins TIA-1 and TIAR link the phosphorylation of eIF-2 alpha to the assembly of mammalian stress granules. J Cell Biol. 1999;147(7):1431–42. PubMed PMID: 10613902.

37. Kedersha N, Cho MR, Li W, Yacono PW, Chen S, Gilks N, et al. Dynamic shuttling of TIA-1 accompanies the recruitment of mRNA to mammalian stress granules. J Cell Biol. 2000;151(6):1257–68. PubMed PMID: 11121440.

38. Skabkin MA, Skabkina OV, Dhote V, Komar AA, Hellen CU, Pestova TV. Activities of Ligatin and MCT-1/DENR in eukaryotic translation initiation and ribosomal recycling. Genes Dev. 2010;24(16):1787–801. doi: 10.1101/gad.1957510. PubMed PMID: 20713520; PubMed Central PMCID: PMCPMC2922506.

39. Dmitriev SE, Terenin IM, Andreev DE, Ivanov PA, Dunaevsky JE, Merrick WC, et al. GTP-independent tRNA delivery to the ribosomal P-site by a novel eukaryotic translation factor. J Biol Chem. 2010;285(35):26779–87. doi: 10.1074/jbc.M110.119693. PubMed PMID: 20566627; PubMed Central PMCID: PMCPMC2930676.

40. Neuvonen M, Kazlauskas A, Martikainen M, Hinkkanen A, Ahola T, Saksela K. SH3 domain-mediated recruitment of host cell amphiphysins by alphavirus nsP3 promotes viral RNA replication. PLoS Pathog. 2011;7(11):e1002383. Epub 2011/11/25. doi: 10.1371/journal.ppat.1002383. PubMed PMID: 22114558; PubMed Central PMCID: PMCPMC3219718.

41 Panas MD, Kedersha N, McInerney GM. Methods for the characterization of stress granules in virus infected cells. Methods. 2015;90:57–64. Epub 2015/04/22. doi: 10.1016/j.ymeth.2015.04.009. PubMed PMID: 25896634.

42. David A, Dolan BP, Hickman HD, Knowlton JJ, Clavarino G, Pierre P, et al. Nuclear translation visualized by ribosome-bound nascent chain puromycylation. J Cell Biol. 2012;197(1):45–57. Epub 2012/04/05. doi: 10.1083/jcb.201112145. PubMed PMID: 22472439; PubMed Central PMCID: PMCPMC3317795.

43. Warner JR, Knopf PM, Rich A. A multiple ribosomal structure in protein synthesis. Proc Natl Acad Sci U S A. 1963;49:122–9. Epub 1963/01/15. PubMed PMID: 13998950; PubMed Central PMCID: PMCPMC300639.

44. Walsh D, Mohr I. Viral subversion of the host protein synthesis machinery. Nat Rev Microbiol. 2011;9(12):860–75. Epub 2011/10/18. doi: 10.1038/nrmicro2655. PubMed PMID: 22002165.

45. Froshauer S, Kartenbeck J, Helenius A. Alphavirus RNA replicase is located on the cytoplasmic surface of endosomes and lysosomes. J Cell Biol. 1988;107(6 Pt 1):2075–86. Epub 1988/12/01. PubMed PMID: 2904446; PubMed Central PMCID: PMCPMC2115628.

46. Pietila MK, van Hemert MJ, Ahola T. Purification of highly active alphavirus replication complexes demonstrates altered fractionation of multiple cellular membranes. J Virol. 2018. doi: 10.1128/JVI.01852-17. PubMed PMID: 29367248; PubMed Central PMCID: PMCPMC5874421.

47. Albulescu IC, Tas A, Scholte FE, Snijder EJ, van Hemert MJ. An in vitro assay to study chikungunya virus RNA synthesis and the mode of action of inhibitors. J Gen Virol. 2014;95(Pt 12):2683–92. doi: 10.1099/vir.0.069690-0. PubMed PMID: 25135884.

48. Kiledjian M, Dreyfuss G. Primary structure and binding activity of the hnRNP U protein: binding RNA through RGG box. EMBO J. 1992;11(7):2655–64. PubMed PMID: 1628625; PubMed Central PMCID: PMCPMC556741.

49. Thandapani P, O’Connor TR, Bailey TL, Richard S. Defining the RGG/RG motif. Mol Cell. 2013;50(5):613–23. doi: 10.1016/j.molcel.2013.05.021. PubMed PMID: 23746349.

50. Tsai WC, Gayatri S, Reineke LC, Sbardella G, Bedford MT, Lloyd RE. Arginine Demethylation of G3BP1 Promotes Stress Granule Assembly. J Biol Chem. 2016;291(43):22671–85. doi: 10.1074/jbc.M116.739573. PubMed PMID: 27601476; PubMed Central PMCID: PMCPMC5077203.

51. Parker F, Maurier F, Delumeau I, Duchesne M, Faucher D, Debussche L, et al. A ras-GTPase-activating protein SH3-domain-binding protein. Mol Cell Biol. 1996;16(6):2561–9. PubMed PMID: WOS:A1996UL97400003.

52. Tourriere H, Gallouzi IE, Chebli K, Capony JP, Mouaikel J, van der Geer P, et al. RasGAP-associated endoribonuclease G3Bp: selective RNA degradation and phosphorylation-dependent localization. Mol Cell Biol. 2001;21(22):7747–60. Epub 2001/10/18. doi: 10.1128/MCB.21.22.7747-7760.2001. PubMed PMID: 11604510; PubMed Central PMCID: PMCPMC99945.

53. Cobos Jimenez V, Martinez FO, Booiman T, van Dort KA, van de Klundert MA, Gordon S, et al. G3BP1 restricts HIV-1 replication in macrophages and T-cells by sequestering viral RNA. Virology. 2015;486:94–104. Epub 2015/10/04. doi: 10.1016/j.virol.2015.09.007. PubMed PMID: 26432022.

54. Bidet K, Dadlani D, Garcia-Blanco MA. G3BP1, G3BP2 and CAPRIN1 are required for translation of interferon stimulated mRNAs and are targeted by a dengue virus non-coding RNA. PLoS Pathog. 2014;10(7):e1004242. Epub 2014/07/06. doi: 10.1371/journal.ppat.1004242. PubMed PMID: 24992036; PubMed Central PMCID: PMCPMC4081823.

55. Vognsen T, Moller IR, Kristensen O. Crystal structures of the human G3BP1 NTF2-like domain visualize FxFG Nup repeat specificity. PLoS One. 2013;8(12):e80947. Epub 2013/12/11. doi: 10.1371/journal.pone.0080947. PubMed PMID: 24324649; PubMed Central PMCID: PMCPMC3852005.

56. Sjoberg EM, Garoff H. The translation-enhancing region of the Semliki Forest virus subgenome is only functional in the virus-infected cell. J Gen Virol. 1996;77 (Pt 6):1323–7. Epub 1996/06/01. PubMed PMID: 8683222.

57. Frolov I, Schlesinger S. Translation of Sindbis virus mRNA: effects of sequences downstream of the initiating codon. J Virol. 1994;68(12):8111–7. PubMed PMID: 7966601.

58. Ventoso I. Adaptive Changes in Alphavirus mRNA Translation Allowed Colonization of Vertebrate Hosts. Journal of Virology. 2012;86(17):9484–94. doi: 10.1128/Jvi.01114-12. PubMed PMID: WOS:000307627800050.

59. Ohshima D, Arimoto-Matsuzaki K, Tomida T, Takekawa M, Ichikawa K. Spatio-temporal Dynamics and Mechanisms of Stress Granule Assembly. Plos Comput Biol. 2015;11(6). doi: ARTN e1004326 10.1371/journal.pcbi.1004326. PubMed PMID: WOS:000357340100038.

60. Siomi MC, Zhang Y, Siomi H, Dreyfuss G. Specific sequences in the fragile X syndrome protein FMR1 and the FXR proteins mediate their binding to 60S ribosomal subunits and the interactions among them. Mol Cell Biol. 1996;16(7):3825–32. PubMed PMID: WOS:A1996UT08600063.

61. McCormick C, Khaperskyy DA. Translation inhibition and stress granules in the antiviral immune response. Nat Rev Immunol. 2017;17(10):647–60. doi: 10.1038/nri.2017.63. PubMed PMID: 28669985.

62. Ulper L, Sarand I, Rausalu K, Merits A. Construction, properties, and potential application of infectious plasmids containing Semliki Forest virus full-length cDNA with an inserted intron. J Virol Methods. 2008;148(1-2):265–70. doi: 10.1016/j.jviromet.2007.10.007. PubMed PMID: WOS:000254720500035.

63. Tamberg N, Lulla V, Fragkoudis R, Lulla A, Fazakerley JK, Merits A. Insertion of EGFP into the replicase gene of Semliki Forest virus results in a novel, genetically stable marker virus. Journal of General Virology. 2007;88:1225–30. doi: 10.1099/vir.0.82436-0. PubMed PMID: WOS:000245493500017.

64. Fukui Y, Yumura S, Yumura TK. Agar-Overlay Immunofluorescence - High-Resolution Studies of Cytoskeletal Components and Their Changes during Chemotaxis. Method Cell Biol. 1987;28:347–56. PubMed PMID: WOS:A1987J148200019.

65. Carpenter AE, Jones TR, Lamprecht MR, Clarke C, Kang IH, Friman O, et al. CellProfiler: image analysis software for identifying and quantifying cell phenotypes. Genome Biol. 2006;7(10):R100. Epub 2006/11/02. doi: 10.1186/gb-2006-7-10-r100. PubMed PMID: 17076895; PubMed Central PMCID: PMCPMC1794559.

66. Gandin V, Sikstrom K, Alain T, Morita M, McLaughlan S, Larsson O, et al. Polysome fractionation and analysis of mammalian translatomes on a genome-wide scale. J Vis Exp. 2014;(87). Epub 2014/06/05. doi: 10.3791/51455. PubMed PMID: 24893926; PubMed Central PMCID: PMCPMC4189431.

67. Hellstrom K, Vihinen H, Kallio K, Jokitalo E, Ahola T. Correlative light and electron microscopy enables viral replication studies at the ultrastructural level. Methods. 2015;90:49–56. doi: 10.1016/j.ymeth.2015.04.019. PubMed PMID: WOS:000365378600007.

68. Tang Z, Ioja E, Bereczki E, Hultenby K, Li C, Guan Z, et al. mTor mediates tau localization and secretion: Implication for Alzheimer’s disease. Biochim Biophys Acta. 2015;1853(7):1646–57. doi: 10.1016/j.bbamcr.2015.03.003. PubMed PMID: 25791428.

