## Supporting Information for "Separate domains of G3BP promote efficient clustering of alphavirus replication complexes and recruitment of the translation initiation machinery"

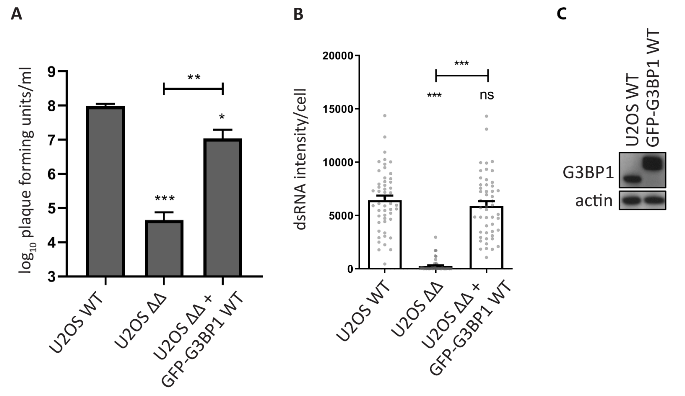


**Supporting Fig S1. GFP-G3BP1 WT rescues SFV replication.** Indicated cells lines were infected with WT SFV at a MOI of 0.1. At 24 hpi, supernatants were collected, and viral titres were quantified by plaque assay on BHK cells. Data are means of three independent experiments. Error bars indicate standard deviation (SD). **B.** Indicated cell lines were infected with WT SFV at MOI 10. At 8 hpi cells were fixed, stained for dsRNA, and dsRNA signal intensities per cell were quantified using CellProfiler software. Bars represent mean + SEM for n = 50 cells per cell line. Individual dsRNA intensities for each analysed cell are shown as grey dots. For statistical analysis indicated cell lines were compared to U2OS WT cells or as indicated. ns P>0.05, *P≤0.05, **P≤0.01, ***P≤0.001. **C.** U2OS WT or U2OS ΔΔ + GFP-G3BP1 WT cells were lysed and analysed by SDS-PAGE and immunoblotting for G3BP1 and actin.

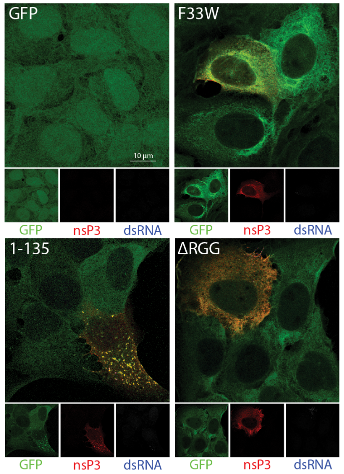


**Supporting Fig S2. CHIKV infection depends on the NTF2-like and RGG domains of G3BP.** Indicated cell lines were infected with CHIKV at MOI 1. At 8 hpi cells were fixed and stained for nsP3 (red) and dsRNA (blue). Representative images of rare dsRNA-positive cells from 3 independent experiments are shown.

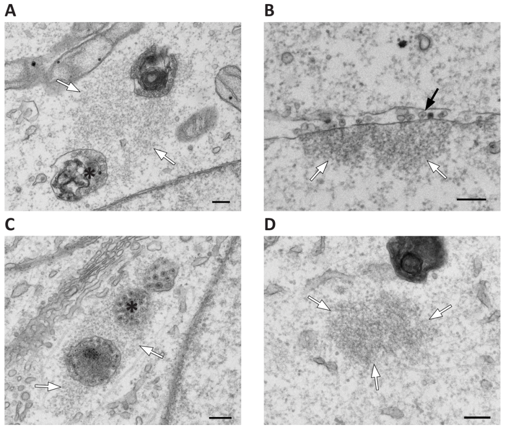


**Supporting Fig S3. Electron-dense patches surround SFV-induced cytopathic vacuoles. A.** U2OS WT cells. **B.** and **C.** U2OS ΔΔ + GFP-G3BP1 WT and **D.** U2OS ΔΔ + GFP-G3BP1 F33W cell lines were infected with WT SFV at MOI 100, fixed at 8 hpi and analysed by transmission electron microscopy. . Black arrows, spherules; White arrows, electron-dense material; Asterisks, cytopathic vacuoles. The scale bar is 200 nm.

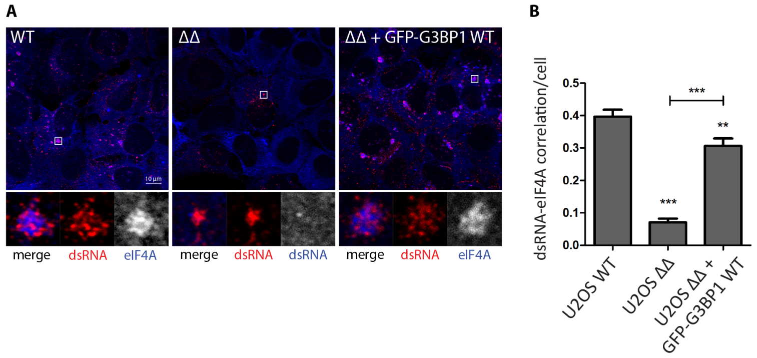


**Supporting Fig S4. Recruitment of eIF4A to SFV CPVs in parental U2OS cells. A.** Indicated cell lines were infected with WT SFV at MOI 10. At 8 hpi cells were fixed and stained for dsRNA (red) and eIF4A (blue/white). Representative images from 2 independent experiments are shown. **B.** Correlation of dsRNA with eIF4A was calculated in CellProfiler based on Pearson's correlation coefficient for n=50 infected cells per sample. Bars represent mean + SEM. For statistical analysis indicated cell lines were compared to U2OS WT cells or as indicated. **P≤0.01, ***P≤0.001.
